# Phage Paride hijacks bacterial stress responses to kill dormant, antibiotic-tolerant cells

**DOI:** 10.1101/2022.01.26.477855

**Authors:** Enea Maffei, Marco Burkolter, Yannik Heyer, Adrian Egli, Urs Jenal, Alexander Harms

## Abstract

Bacteriophages are fierce viral predators with no regard for pathogenicity or antibiotic resistance of their bacterial hosts^1,2^. Despite early recognition of their therapeutic potential and the current escalation of bacterial multidrug resistance, phages have so far failed to become a regular treatment option in clinical practice^3-5^. One reason is the occasional discrepancy between poor performance of selected phages *in vivo* despite high potency *in vitro*^3,4,6,7^. Similar resilience of supposedly drug-sensitive bacterial infections to antibiotic treatment has been linked to persistence of dormant cells inside patients^8,9^. Given the abundance of non-growing bacteria also in the environment^10,11^, we wondered whether some phages can infect and kill these antibiotic-tolerant cells. As shown previously^12-16^, most phages failed to replicate on dormant hosts and instead entered a state of hibernation or pseudolysogeny. However, we isolated a new *Pseudomonas aeruginosa* phage named Paride with the exciting ability to directly kill dormant, antibiotic-tolerant hosts by lytic replication, causing sterilization of deep-dormant cultures in synergy with the β-lactam meropenem. Intriguingly, efficient replication of Paride on dormant hosts depends on the same bacterial stress responses that also drive antibiotic tolerance. We therefore suggest that Paride hijacks weak spots in the dormant physiology of antibiotic-tolerant bacteria that could be exploited as Achilles’ heels for the development of new treatments targeting resilient bacterial infections.

## Introduction

Bacteriophages (or short “phages”) are the viruses infecting bacteria and act as top predators of the microbial world^1,2^. It is therefore intuitive to use phages as powerful allies in the fight against bacterial infections, and they have been applied to treat patients for over a century^3,4^. However, a lack of reliability and technical difficulties caused rapid disappearance of phage therapy in the West once antibiotic drugs became available^3,4^. This trend has reversed in recent years when phages, owing to advances in medicine and biotechnology, have been increasingly used as attractive treatment options for multidrug-resistant bacterial infections^3,4,17-19^. Given the bleak prospects of the antibiotic resistance crisis with around five million deaths linked to resistant bacterial infections in 2019^5^, this trend is likely going to continue.

However, despite justified fears of an imminent post-antibiotic “era of untreatable infection”^20^, antibiotic resistance is not the only threat to patients suffering from bacterial infections. Even without genetic resistance, bacteria can defy antimicrobial treatment in stressed and starved dormant states that are common in nature and in patients during chronic or relapsing infections^10,11^. This antibiotic tolerance causes a slower killing of a genetically sensitive bacterial population under drug treatment and is often accompanied by the formation of deep-dormant, highly antibiotic-tolerant cells known as bacterial persisters^8,9,21^. Despite decades of intensive research, the molecular mechanisms underlying antibiotic tolerance and bacterial persisters are still hotly debated and no effective treatments are available^8,9,21^. However, it seems clear that antibiotic tolerance is linked to a dormant physiology of bacterial pathogens in which the cellular processes commonly poisoned by bactericidal antimicrobials are tuned down or inactive^8,9,21^.

While some cases of successful phage therapy have targeted chronic bacterial infections that were highly resilient to antibiotic treatment^17-19^, it has been noted since long that in other clinical cases “the bacteriophage, which acts so well *in vitro*, does not have a similar action *in vivo*”^7^. Consequently, it is increasingly recognized that the bacterial physiology at the infection site is a key parameter for the success of phage therapy and can greatly influence the infectivity of these viruses^6,22^. We therefore wondered if bacteriophages can infect dormant, antibiotic-tolerant bacteria that are likely abundant in their natural habitats and whether this ability differs between phages.

Previous work suggested that the productivity of phage infections is positively correlated with host growth rate and that fully growth-arrested cells are refractory to phage replication^12,23-26^. The resource limitation during bacterial dormancy therefore seems to be a major barrier to phage infection, and this phenomenon is weaponized against these viruses by a wide variety of bacterial immunity systems that shut down cellular physiology upon phage infection to restrict viral spread^27,28^. Consequently, commonly studied obligately virulent phages either avoid adsorption to dormant bacteria^29^ or hibernate in the low-energy physiology of these cells until nutrients become available again and lytic replication resumes^12-15^, a phenomenon also known as pseudolysogeny^16^. True lysogeny is restricted to so-called temperate phages which can integrate their genome into the host’s genome to form a transient evolutionary coalition, a decision that is heavily skewed towards lysogeny in starved host cells^30^. Direct lytic replication of bacteriophages on dormant hosts has been reported only rarely^31,32^, but bears a great potential to improve phage therapy and to inspire the development of better treatment options for chronic bacterial infections.

In this study we therefore performed large-scale bacteriophage isolation experiments to isolate new phages with the ability to directly kill antibiotic-tolerant, dormant cells of *Escherichia coli* or *Pseudomonas aeruginosa* by lytic replication. While most phages seemed to hibernate in these hosts with variable success, we isolated a new *P. aeruginosa* phage named Paride that uniquely replicates on deep stationary-phase cultures of laboratory and clinical strains of this organism. Intriguingly, we found that Paride can even sterilize deep-stationary phase cultures of *P. aeruginosa* via phage-antibiotic synergy with the β-lactam meropenem. Intriguingly, the replication of Paride on dormant hosts largely depended on the bacterial starvation and stress response signalling that is also required for the antibiotic tolerance of these bacteria. This suggests that Paride specifically exploits weak spots in the resilient physiology of dormant bacteria that could be targeted as Achilles’ heels by new treatment options.

## Results

### Commonly studied bacteriophages cannot replicate on deep-dormant *E. coli* and *P. aeruginosa*

We initially explored whether a wide range of different phages including commonly used laboratory models can kill deep-dormant cultures of *Escherichia coli* or *P. aeruginosa* by direct replication. Given that well-chosen and strictly controlled assay conditions are crucial for meaningful experiments with dormant bacteria^9,21,33,34^, we had previously established a rigorous methodology to study antibiotic tolerance of regularly growing and deep-dormant cultures of these organisms^35^. Briefly, bacteria are grown in a fully defined growth medium and then challenged with antibiotics or, in the current study, bacteriophages during exponential growth or after close to 48h in stationary phase. Under these conditions, fast-growing cultures of both bacterial species are readily cleared by antibiotic treatment and highly permissive to replication by all tested bacteriophages (Extended Data Fig. 1a,b). Conversely, the deep-dormant cultures displayed massive antibiotic tolerance and did not allow replication of any of the tested bacteriophages (Fig. 1a-e). Instead, most phages rapidly adsorbed and then seemed to enter a state of hibernation in dormant hosts that is apparent as a stable number of infected cells over time and had previously already been observed with *E. coli* phage T4 and *P. aeruginosa* phage UT1^14,15^.

**Figure 1:**
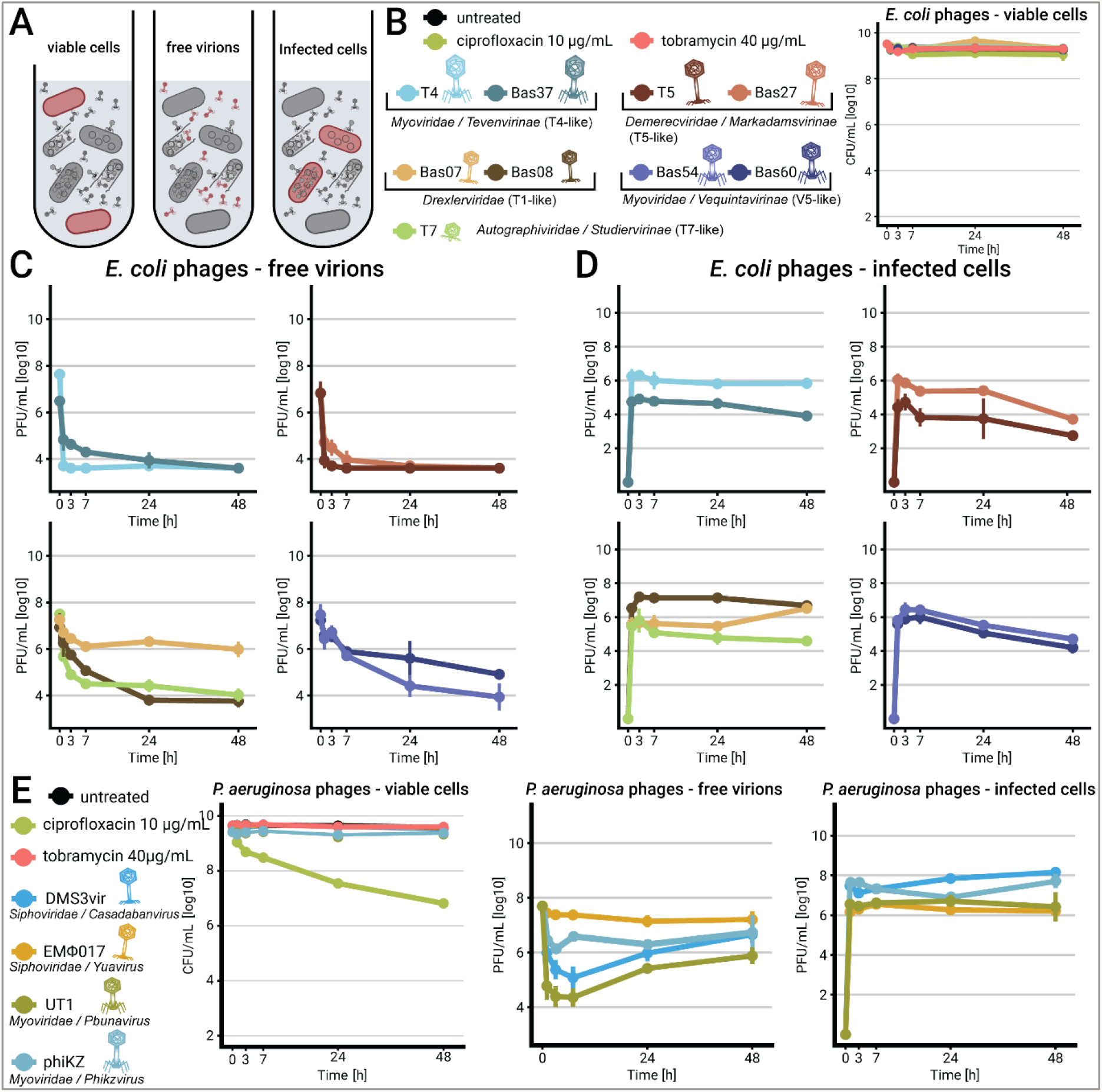
Most bacteriophages cannot replicate on deep-dormant *E. coli* or *P. aeruginosa*. (**a**) Schematic of metrics that were recorded during phage infection experiments. (**b-e**) Deep-dormant cultures of *E. coli* K-12 MG1655 or *P. aeruginosa* PAO1 *Δpel Δpsl* were treated with antibiotics or phages (multiplicity of infection (MOI) ≈ 0.01) and viable colony forming units (CFU/ml) as well as plaque-forming units (PFU/ml) of free phages and infected cells were recorded over time. Data points and error bars show average and standard error of the mean for 2-3 independent experiments. Limits of detection are 2 log10 CFU/mL for viable cells, 3.6 log10 PFU/mL for free phages and 2.6 log10 PFU/mL for infected cells.

### Bacteriophage Paride can kill deep-dormant *P. aeruginosa* by direct lytic replication

We therefore adapted our previously developed bacteriophage isolation setup^35^ to broadly screen for new phages that could directly replicate on dormant hosts by using deep-dormant cultures of *E. coli* or *P. aeruginosa* as bait (Fig. 2a). These experiments resulted in the isolation of bacteriophage Paride, a *P. aeruginosa* phage that rapidly adsorbs to deep-dormant host cells and then massively replicates, killing >99% of the bacterial population and causing the culture to lyse (Fig. 2b and Extended Data Fig. 2a,b). Paride forms virions of myovirus morphotype and has a large genome of 287’267 bp, i.e., far beyond the 200kb threshold defining “jumbo phages” (*NCBI GenBank accession not yet available*; Fig. 2c)^36^. Phylogenetic analyses revealed that Paride is a close relative of previously described phages PA5oct and MIJ3 (Fig. 2d)^37,38^. However, Paride is not noticeably related to well-studied *P. aeruginosa* jumbo phage phiKZ which famously forms a “phage nucleus” in infected cells^39^, cannot replicate on dormant hosts (Fig. 1e), and belongs to an entirely different clade of jumbo phages^40^.

**Figure 2:**
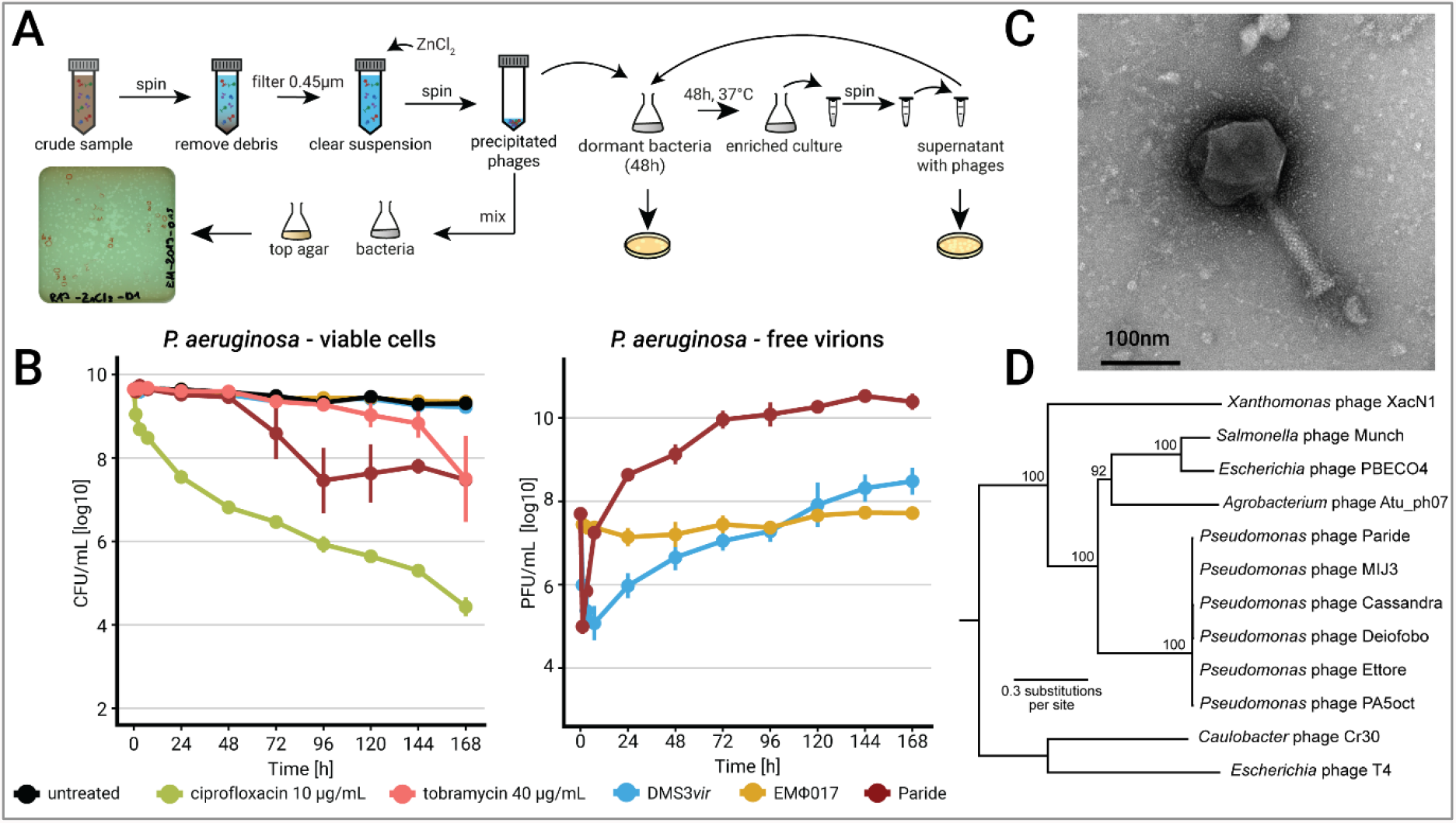
Paride is a new *P. aeruginosa* phage that can replicate on deep-dormant host cells. (**a**) Enrichment strategy to isolate phages that can replicate on stationary phase cultures (see *Methods* for details). (**b**) Deep-dormant cultures of *P. aeruginosa* were treated with antibiotics or phages (MOI ≈ 0.01) and viable CFU/ml as well as free phages were recorded over time. Data points and error bars show the average of three biological replicates and standard error of the mean. Limits of detection are 2 log10 CFU/mL for viable cells and 3.6 log10 PFU/mL for free phages. (**c**) Negative stain TEM micrograph of phage Paride. (**d**) Maximum-likelihood phylogeny of Paride and other group 2.2 jumbo phages^40^ with phages T4 and Cr30 as outgroup (see *Methods*). Bootstrap support is shown if >70.

Repeated attempts at isolating different phages that can replicate on dormant, antibiotic-tolerant bacteria exclusively uncovered diverse close relatives of Paride that we called Cassandra, Deiofobo, and Ettore (Fig. 2d and Supplementary Table 1) but no other phage, suggesting that this ability is very rare. We therefore explored whether the observed replication of Paride on deep-dormant cultures might be a laboratory artifact from the combination of this phage and the *P. aeruginosa* PAO1 model strain. However, Paride also readily replicated on stationary-phase cultures of different susceptible *P. aeruginosa* strains from a collection of clinical isolates (Fig. 3a and Extended Data Fig. 3a), demonstrating that this phenomenon is not restricted to the PAO1 laboratory strain.

**Figure 3:**
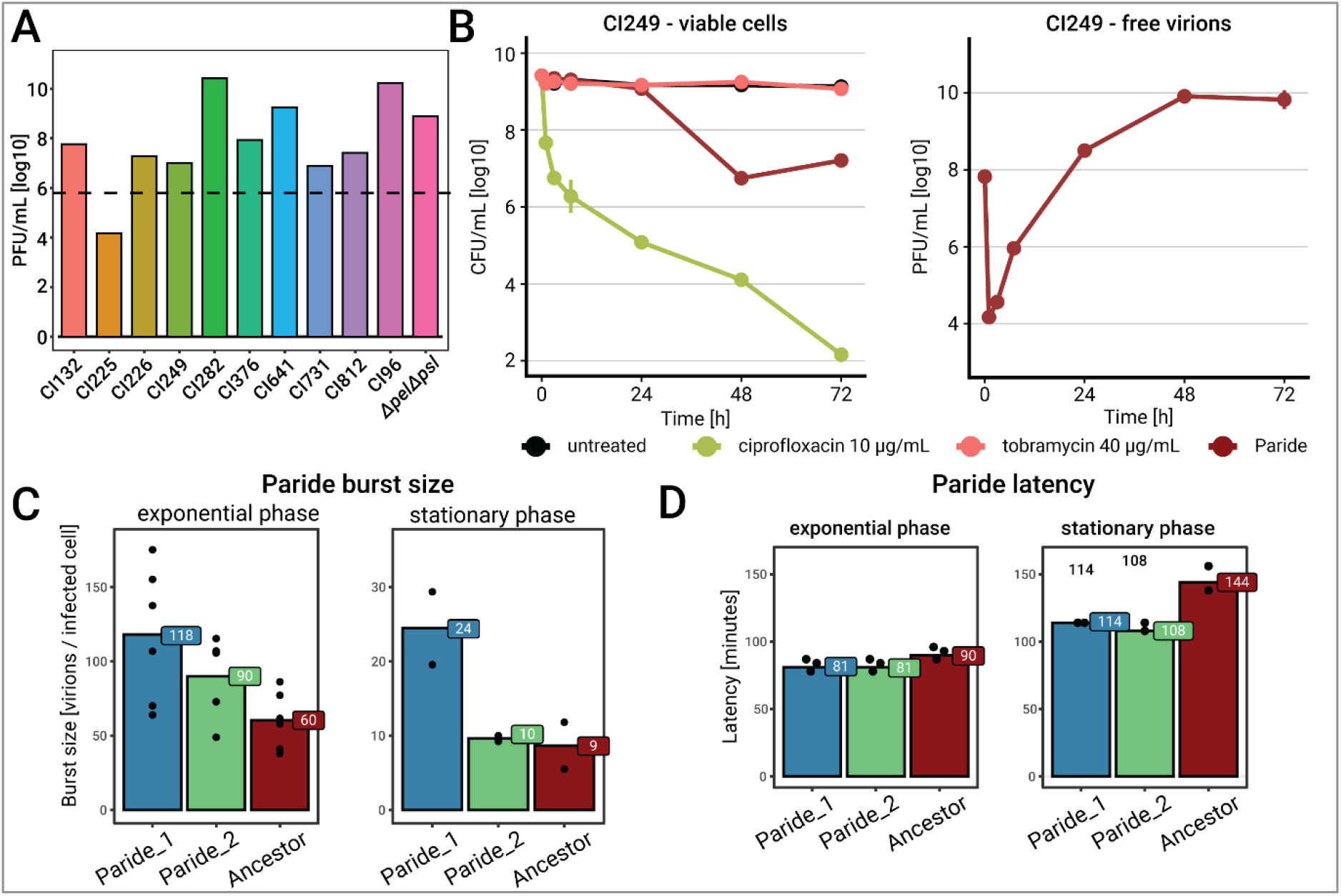
Infection of clinical *P. aeruginosa* isolates by Paride and one-step growth curves. (**a**) Free phage titers of Paride after infecting deep-dormant cultures of different clinical isolates of *P. aeruginosa* for 48h (dashed line: inoculum). (**b**) Deep-dormant cultures of *P. aeruginosa* clinical isolate CI249 were treated with antibiotics or Paride (MOI ≈ 0.01) and viable CFU/ml as well as free phages were recorded over time. Due to lack of robust growth in M9Glc this experiment was performed in M9Rich (see Extended Data Fig. 3a for a control experiment with the *P. aeruginosa* PAO1 *Δpel Δpsl* laboratory wildtype in this medium). Data points and error bars show average and standard error of the mean of three independent experiments. Limits of detection are 2 log10 CFU/mL for viable cells, 3.6 log10 PFU/mL for free phages. (**c**,**d**) Burst size and latency of ancestral Paride as well as two evolved clones passaged on deep-dormant cultures were determined by one-step growth experiments (see also Extended Data Fig. 3b,c). Data bars represent the average of 2-6 independent experiments and all individual data points are shown.

### Quantitative assessment of Paride infections and experimental evolution

We then performed one-step growth experiments to quantify the speed and productivity of Paride infections. For regularly growing hosts under our experimental conditions, we determined a burst size of around 60 (i.e., virions produced per infected cell, Fig. 3c) and a latency period of 1.5h (i.e., infection time needed to generate new virions; Fig. 3d). When infecting dormant hosts, Paride showed a reduced burst size of ca. 9 and a prolonged latency period of ca. 2.5h (Fig. 3c,d). With view to possible medical relevance of Paride’s ability to replicate on dormant hosts, we performed experimental evolution to improve this ability by serially passaging two independent lines on deep-stationary phase cultures for around 600 generations. Both lines improved burst size and latency, though the improvement of burst size was more pronounced in the Paride_1 lineage while the improvement of latency was more pronounced in the Paride_2 lineage (Fig. 3c,d). This result suggests that the two lineages improved overall infection efficiency by convergent evolution via different routes. None of these improvements was specific to infecting deep-dormant cultures, i.e., experimental evolution had generally improved Paride infectivity.

### Paride targets the outer core of *P. aeruginosa* LPS as essential host receptor

Phage PA5oct had previously been shown to have a partial requirement for type IV pili and lipopolysaccharides (LPS) of its *P. aeruginosa* host^41^ which are both very common – though usually distinct – phage receptors on this organism^42^. We readily confirmed that Paride infectivity is partially compromised in absence of either type IV pili (Δ*pilA*) or O-antigen (Δ*wbpL*; Fig. 4a,b and Extended Data Fig. 4). Using a panel of spontaneously resistant mutants, we determined that hosts with deeper truncations of the LPS core below the O-antigen (Δ*galU* or Δ*ssg*) are completely and not only partially resistant to Paride (Fig. 4a,b). These genes had already previously been implicated in resistance to LPS-targeting phages^43-46^. Based on these results, we conclude that the essential terminal receptor for Paride infections is located in the outer core of the *P. aeruginosa* LPS and probably includes its αα-glucose(III) moiety (Fig. 4a,b and Extended Data Fig. 4; see also in *Methods* and Supplementary Table 2)^47-49^.

**Figure 4:**
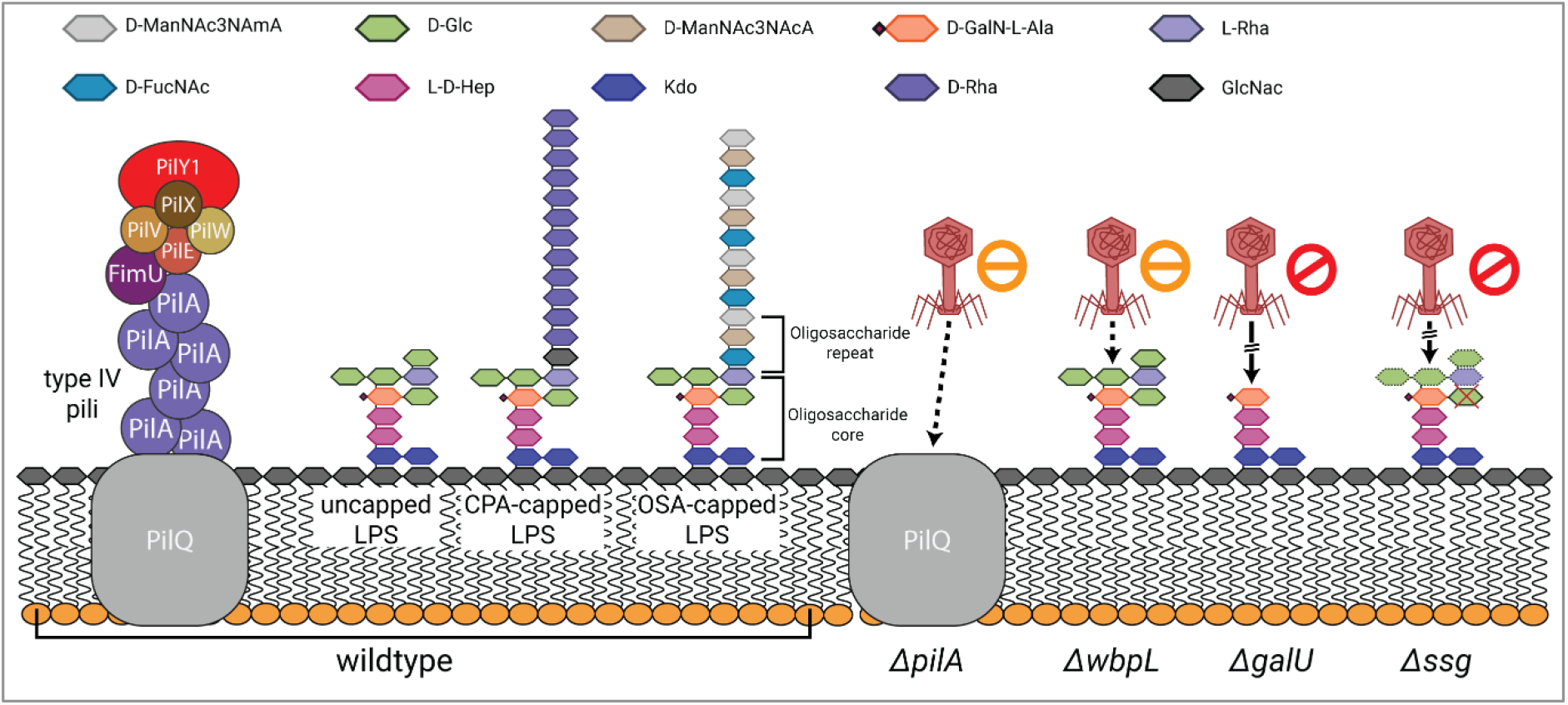
Paride binds the outer LPS core of *P. aeruginosa* as essential host receptor. Schematic representation of the P. aeruginosa PAO1 Δpel Δpsl cell surface and susceptibility of different mutants to Paride in top agar assays (see Methods for details). Top agar assays underlying the interpretation shown in this illustration are shown in Extended Data Fig. 4. CPA = common polysaccharide antigen, OSA = O-specific antigen

### Productive infection of deep-dormant hosts by Paride requires functional stress responses

Given the correlation of antibiotic tolerance and resilience to phage infections for deep-dormant bacteria (compare Fig. 1 and Extended Data Fig. 1)^35^, we hypothesized that the bacterial core signalling orchestrating their dormant physiology might be responsible for both phenomena. In many Gram-negatives, the stringent response second messenger (p)ppGpp and stress response sigma factor RpoS together tune down cellular processes in stationary phase which is thought to cause antibiotic tolerance^9,50-52^. We therefore tested whether knocking out the makers and breakers of (p)ppGpp (*relA* and *spoT*) or *rpoS* might sensitize non-growing *P. aeruginosa* to phages other than Paride. As expected, the *ΔrpoS* and *ΔrelA ΔspoT* mutants displayed greatly reduced antibiotic tolerance under all tested conditions (Fig. 5a,b and Extended Data Fig. 5a-d). However, they were still not more permissive to infection by control phages when grown into growth arrest and instead were highly refractory to infection by Paride under these conditions (Fig. 5a,b and Extended Data Fig. 5a,b). Notably, Paride infections of regularly growing *ΔrpoS* or *ΔrelA ΔspoT* strains were indistinguishable from the parental wildtype (Extended Data Fig. 5c,d). These results suggest that the ability of Paride to directly replicate on non-growing hosts depends on subversion of the regular stationary phase physiology of deep-dormant hosts.

**Figure 5:**
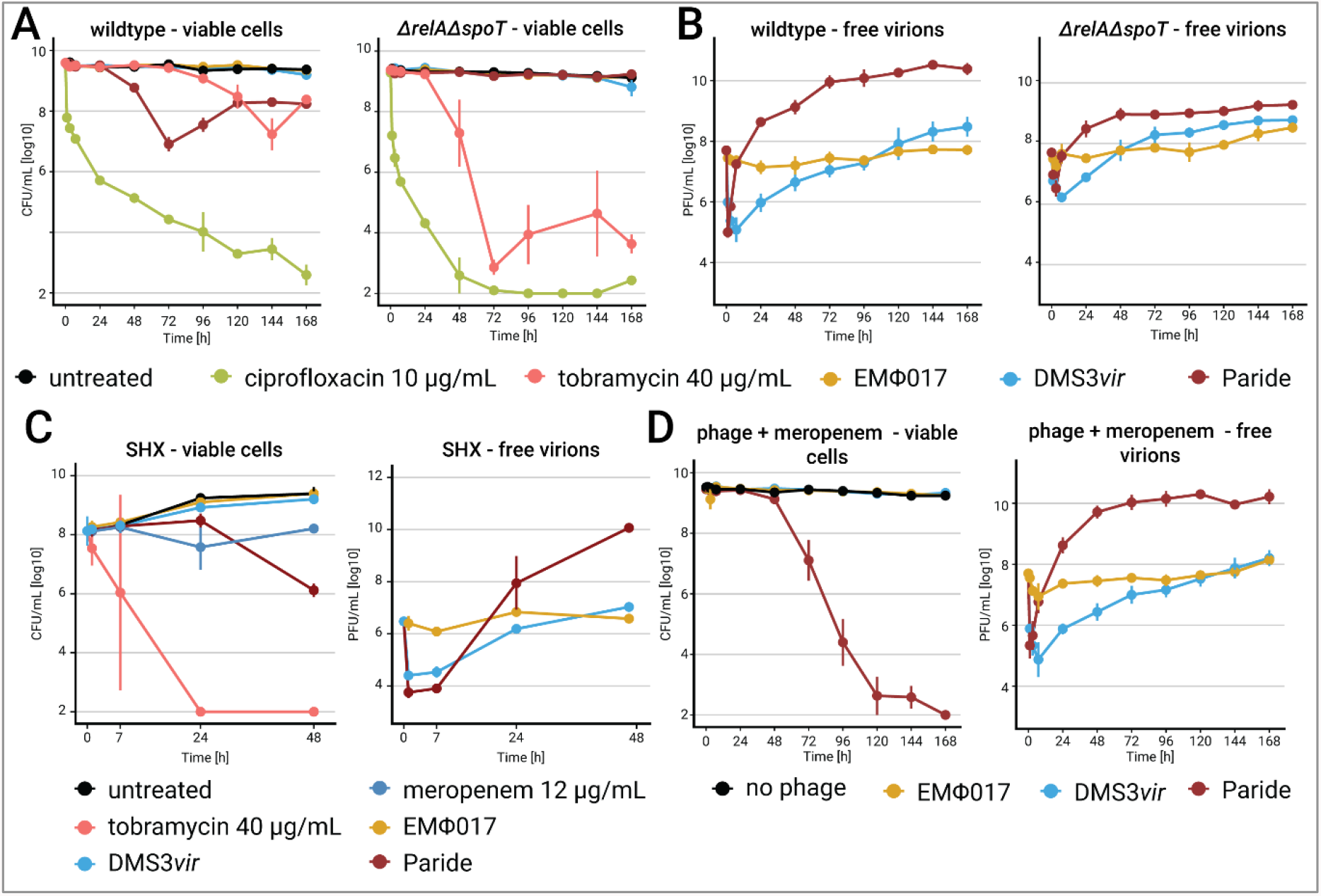
Paride requires stress responses for infections of dormant hosts and shows phage-antibiotic synergy with meropenem. (**a**,**b**) Deep-dormant cultures of *P. aeruginosa* PAO1 *Δpel Δpsl* (wildtype) and its *ΔrelA ΔspoT* derivative both grown in M9Rich were treated with antibiotics or phages (MOI ≈ 0.01) and viable CFU/ml as well as free phages were recorded over time. (**c**) Growing cultures of *P. aeruginosa* were treated with 1 mM of DL-serine hydroxamate for 12 hours and then challenged with antibiotics or phages (MOI ≈ 0.01). Viable CFU/ml as well as free phages were recorded over time (see Extended Data Fig. 5e for a control experiment without SHX). (**d**) Deep-dormant cultures of *P. aeruginosa* were treated with meropenem in combination with phages (MOI ≈ 0.01) and viable CFU/ml as well as free phages were recorded over time. Analogous experiments with ciprofloxacin and tobramycin are shown in Extended Data Fig. 5f,e. Data points and error bars show the average of three (d: two) independent experiments and their standard error of the mean. Limits of detection are 2 log10 CFU/mL for viable cells and 3.6 log10 PFU/mL for free phages.

As an independent control for these experiments and the deep-dormant cell infections in general, we treated regularly growing *P. aeruginosa* cultures with the stringent response inducer serine hydroxamate (SHX) to induce strong starvation-like signalling (Fig. 5c)^53^. Similar to deep dormancy, SHX pre-treatment induced considerable antibiotic tolerance and resilience to phage infection, while Paride maintained its ability to replicate under these conditions (Fig. 5c and Extended Data Fig. 5e).

### Phage-antibiotic synergy of Paride and meropenem sterilizes deep-dormant cultures

The combined treatment of bacterial infections with antibiotic drugs and bacteriophages can have a strong synergistic effect, but these interactions are difficult to predict and mostly applied empirically^54^. We therefore investigated whether the combination of Paride with antibiotic drugs might enable the killing of more than the ca. 99% of deep-dormant cells that are eliminated by the phage alone before a plateau of phenotypic resistance is reached (see Fig. 2b). For this purpose, the Paride infection experiments of deep-dormant cultures were repeated in combination with lethal concentrations of the fluoroquinolone ciprofloxacin, the aminoglycoside tobramycin, or the β-lactam meropenem. Under our experimental conditions, combining phage infections with ciprofloxacin or tobramycin had no effect beyond the bactericidal action of the antibiotics alone (Extended Data Fig. 5f,g). However, treatment with meropenem and Paride together resulted in complete sterilization of deep-dormant *P. aeruginosa* cultures to the detection limit (Fig. 4e) even though meropenem alone had no detectable effect under these conditions even during prolonged treatment^35^.

## Discussion

### Bacteriophages use different strategies to infect dormant host cells

Our study confirms the previous notion that bacteriophages are generally unable to directly replicate on dormant bacteria^12,26-28^, but presents a new *P. aeruginosa* phage named Paride with the unique ability to kill deep-dormant bacteria by direct lytic replication in our infection model (Figs. 1 and 2b). Interestingly, efficient replication of Paride on growth-arrested hosts specifically requires cellular stress responses in form of (p)ppGpp and RpoS signalling that are dispensable for infections of growing hosts (Figs. 5a,b and Extended Data Fig. 5a-d). These results indicate that Paride subverts certain aspects of the host’s dormant physiology to enable direct replication, e.g., by mobilizing resources and energy that are stored away in regular stationary phase cells and might not be available in growth-arrested hosts lacking the core stress and starvation responses^51,52^. It will be interesting to see future work studying the molecular details of how Paride overcomes the resilient physiology of dormant host cells. As a jumbo phage with a large genome of almost 300kb^36,40^, Paride could easily encode a dedicated genetic program for this purpose analogous to, e.g., the fascinating nucleus-like structure assembled by jumbo phage phiKZ to protect its genome from bacterial immunity^39^.

Unlike Paride, most other phages enter a state of hibernation in deep-dormant host cells that is very stable in some cases (e.g., T4, Bas08, UT1, phiKZ) and less so in others (Fig. 1d,e). Hibernation of phage T4 hinges on the arrest of its lytic program in dormant cells after degrading the host chromosome and in dependence on lysis control gene *rI*^14,55^. This would suggest that hibernation can be a phage-imposed strategy activated after irreversible takeover of a dormant cell to postpone replication until more resources are available to maximize viral productivity. It is tempting to speculate that hibernation truly represents a form of “pseudolysogeny” in which the virus seeks shelter from UV radiation and other environmental hazards inside bacterial cells^16^. It will be interesting to further investigate if phage hibernation represents a general viral strategy or if it is simply caused by viral paralysis in resource-limited dormant hosts, e.g., as a kind of physiological defence against phage infection. Notably, recent work supported this hypothesis by demonstrating that host dormancy strongly promotes the acquisition of CRISPR-Cas immunity against infecting phages^56^.

Another phenomenon observed with deep-dormant cultures is that some *P. aeruginosa* phages seem to rapidly adsorb and enter hibernation after which the number of free virions increases again over the next days, but never above the inoculum (Fig. 1e). In the absence of evidence for viral replication or bacterial killing, it seems evident that this phenomenon is caused by reversible adsorption of a minor subpopulation of phages that fail to irreversibly bind the terminal surface receptor and inject their genome. This would be reminiscent of *E. coli* phages T1 or phi80 that can only irreversibly bind at their FhuA receptor and inject their genome when FhuA is energized by the TonB system^29^. However, the different phages showing this behaviour in our study bind to different receptors on the cell surface with phiKZ and DMS3*vir* targeting type IV pili while PB1-like phages like UT1 generally target the *P. aeruginosa* LPS^41^. Thus, this phenomenon could instead be caused by phenotypic resistance of a bacterial subpopulation (similar to the ca. 1% of cells resisting Paride infections in stationary phase, Fig. 2b) or phenotypic heterogeneity on the phage side as observed previously^57^.

### Different results obtained with different experimental models for “stationary phase”

This work was largely performed using our previously described experimental setup for studying the biology of deep-dormant bacteria that is based on defined culture media and rigorously controlled assay conditions^35^. It is important to highlight the technical details of these assays because experimentation with antibiotic-tolerant, dormant bacteria is highly sensitive to seemingly small changes in the assay setup^9,21,33,34^. The “stationary phase” cells in our experiments have been in a non-growing state under severe nutrient limitation for almost 48h. They are stably dormant without loss of cell viability and are non-dividing because we do not observe significant killing even under prolonged treatment with lethal concentrations of β-lactam drugs that poison bacterial cell wall biosynthesis, eliminating any intermittently growing cell (Fig. 2b)^35,58^. This is different from other methodologies sometimes used in the field. In particular, cultures grown in LB broth show significant spontaneous cell death, an equilibrium of growing and dying cells, and considerable killing by β-lactams^59,60^.

In our controlled assay setup we could reproduce some previously published results such as, e.g., the hibernation of *E. coli* phage T4 and *P. aeruginosa* phage UT1^14,15^ in dormant hosts (Fig. 1d,e). However, we failed to observe an ability of phages T7 and UT1 to replicate on stationary phase cells that had been reported previously^31,32^. It seems likely to us that this difference is due to differences in the “stationary phase” physiology of the bacteria that have been investigated because the cells in our assay setup might be particularly dormant as evidenced, e.g., by their strong antibiotic tolerance^35^. Notably, phage T7 has previously been reported to be highly proficient at infecting highly starved and stressed (albeit growing) cells^61^, making it plausible that this phage may be able to replicate on non-growing cells at a less dormant stage of stationary phase.

### Sterilization of deep-dormant cultures by phage-antibiotic synergy of Paride and meropenem

One of the most exciting results of our study is that a combination of Paride and meropenem can sterilize deep-dormant cultures to the extent that no colony-forming units can be recovered anymore (Fig. 5d, at least 7 logs reduction of bacterial counts). Notably, Paride alone can kill around 99% of cells in these stationary phase cultures (Fig. 2b) while meropenem alone or in combination with other phages is completely ineffective^35^. We interpret this phage-antibiotic synergy as a chain reaction initiated by lysis of some deep-dormant cells by Paride. Molecules released from these cells might cause resuscitation of phenotypically phage-resistant bystanders and enable their effective killing by meropenem as soon as cell wall biosynthesis resumes and can be poisoned by β-lactams^58^. While it is intuitive that resuscitation might be caused by nutrients, it could also be due to cell wall fragments serving as a resuscitation signal for dormant cells^62^ that is produced by Paride infections either accidentally or via an evolved phage-controlled mechanism. Understanding this phage-antibiotic synergy on the molecular level might enable us to design new treatment options for resilient bacterial infections based on the forced resuscitation of deep-dormant, drug-tolerant bacteria.

## Extended Data Figure legends

Extended Data Fig.1: Time-kill curves of regularly growing *E. coli* and *P. aeruginosa*

(**a**) Fast-growing cultures of *E. coli* were treated with antibiotics or phages (MOI ≈ 0.001) and viable CFU/ml as well as free phages were recorded over time. (**b**) Fast-growing cultures of *P. aeruginosa* were treated with antibiotics or phages (MOI ≈ 0.001) and viable CFU/ml as well as free phages were recorded over time. Data points represent the average of 2-3 independent experiment and error bars show their standard error of the mean. Limits of detection are 2 log10 CFU/mL for viable cells, 3.6 log10 PFU/mL for free phages and 2.6 log10 PFU/mL for infected cells.

Extended Data Fig. 2: Infection of regularly growing *P. aeruginosa* with Paride and lysis of deep-dormant cultures by the same phage

(**a**) Regularly growing cultures of *P. aeruginosa* were treated with antibiotics or phages (MOI ≈ 0.001) and viable CFU/ml as well as free phages were recorded over time. Data points represent the average of three independent experiment and error bars show their standard error of the mean. Limits of detection are 2 log10 CFU/mL for viable cells, 3.6 log10 PFU/mL for free phages and 2.6 log10 PFU/mL for infected cells. (**b**) Representative picture of a typical, deep-dormant *P. aeruginosa* culture at 96h post treatment start in M9Glc, untreated (left) and infected with Paride (right).

Extended Data Fig. 3: – Stationary phase time-kill curves of *P. aeruginosa* in M9Rich and one-step growth curves of Paride

(**a**) Deep-dormant cultures of *P. aeruginosa* grown in M9Rich were treated with antibiotics or phages (MOI ≈ 0.01) and viable CFU/ml as well as free phages were recorded over time. Data points represent the average of three independent experiment and error bars show their standard error of the mean. Limits of detection are 2 log10 CFU/mL for viable cells, 3.6 log10 PFU/mL for free phages and 2.6 log10 PFU/mL for infected cells. (**b, c**) Free virions of one-step growth experiments with ancestral Paride and two lineages evolved on deep-dormant cultures for ca. 600 generations were recorded over time in fast-growing cultures (b) and stationary phase cultures (c). Data points represent the average of six (regularly growing) and two (stationary phase) independent experiments, respectively. The dashed line represents the limit of detection (3.6 log10 PFU/mL).

Extended Data Fig. 4: – Top agar assays with Paride and control phages on different surface receptor mutants of *P. aeruginosa* PAO1

Top agars were set up with *P. aeruginosa* PAO1 *Δpel Δpsl* (wildtype) and different constructed or spontaneously isolated mutants lacking functional expression of one or more surface receptor genes before infection with serial dilutions of phage Paride and control phages E79 (targeting the LPS core^63,64^), newly isolated phage EMΦ31 (targeting the LPS O-antigen), and DMS3*vir* (targeting type IV pili^65^). Arrows highlight opaque plaque formation of phage Paride on several mutants. Strain EM-307 is a spontaneously isolated “brown mutant” as described previously with a large deletion around *galU*^43^. The data are summarized in Supplementary Table 2.

Extended Data Fig. 5: Additional data regarding Paride infections of Δ*rpoS* and Δ*relA* Δ*spot* mutants, in presence of SHX, and phage-antibiotic combination treatments

(**a**) Deep-dormant cultures of *P. aeruginosa* PAO1 *Δpel Δpsl* (wildtype) and its *ΔrpoS* derivative both grown in M9Glc were treated with antibiotics or phages (MOI ≈ 0.01) and viable CFU/ml as well as free phages were recorded over time. (**b**) Fast-growing cultures of *P. aeruginosa* PAO1 *Δpel Δpsl* (wildtype) and its *ΔrpoS* derivative both grown in M9Glc were treated with antibiotics or phages (MOI ≈ 0.001) and viable CFU/ml as well as free phages were recorded over time. (**c**,**d**) Fast-growing cultures of *P. aeruginosa* PAO1 *Δpel Δpsl* (wildtype) and its *ΔrelA ΔspoT* derivative both grown in M9Rich were treated with antibiotics or phages (MOI ≈ 0.001) and viable CFU/ml as well as free phages were recorded over time. (**e**) Growing cultures of *P. aeruginosa* were processed as described for the SHX treatment of Fig. 5b (just as a control without SHX) and then challenged with antibiotics or phages (MOI ≈ 0.01). Viable CFU/ml as well as free phages were recorded over time. (**f, g**) Deep-dormant cultures of *P. aeruginosa* were treated with tobramycin (f) or ciprofloxacin (g) in combination with phages (MOI ≈ 0.01) and viable CFU/ml as well as free phages were recorded over time. Data points and error bars show the average of three (e: two) independent experiments and their standard error of the mean. Limits of detection are 2 log10 CFU/mL for viable cells and 3.6 log10 PFU/mL for free phages.

## Methods

### Preparation of culture media and solutions

Lysogeny Broth (LB) was prepared by dissolving 10 g/L tryptone, 5 g/L yeast extract, and 10 g/L sodium chloride in Milli-Q H_2_O and sterilized by autoclaving. LB agar plates were prepared by supplementing LB medium with agar at 1.5% w/v before autoclaving. M9Glc was prepared as described previously^35^. The M9Rich culture medium was conceived as a variant of regular M9Glc medium supplemented with 10% v/v LB medium (prepared without NaCl) to promote the growth of diverse strains^35^. It was prepared from sterilized components by mixing (for 50 mL) 33.75 mL Milli-Q H2O, 10 mL 5× M9 salts solution, 5 mL LB medium without NaCl, 500 μl 40% w/v D-glucose solution, 100 μL 1 M MgSO4, and 5 μL 1 M CaCl2 using sterile technique. Unless indicated otherwise, all components were sterilized by filtration (0.22 μm). Phosphate-buffered saline (PBS) was prepared as a solution containing 8 g/L NaCl, 0.2 g/L KCl, 1.44 g/L Na_2_HPO_4_x2H_2_O, and 0.24 g/L KH_2_PO_4_ with the pH adjusted to 7.4 using 10 M NaOH and sterilized by autoclaving. SM buffer was prepared as 0.1M NaCl, 10mM MgSO_4_, and 0.05M Tris (pH 7.5) using sterile technique.

### Bacterial handling and culturing

*E. coli* and *P. aeruginosa* strains were routinely cultured in LB medium at 37°C in glass culture tubes or Erlenmeyer flasks with agitation at 170 rpm. For all antibiotic treatment and phage infections assays, the bacteria were instead grown in M9Glc or M9Rich. Clinical isolates of *P. aeruginosa* often showed fastidious growth requirements and were always cultivated in M9Rich. LB agar plates were routinely used as solid medium. Selection for genetic modifications or plasmid maintenance was performed with gentamicin at 20 μg/mL, ampicillin 100 μg/mL, oxytetracycline 12.5 μg/mL for *E. coli* or gentamicin at 30 μg/mL, carbenicillin 100 μg/mL, and oxytetracycline 100 μg/mL for *P. aeruginosa*.

### Bacteriophage handling and culturing

Bacteriophages (listed in Supplementary Table 1) were generally cultured using the double-agar overlay (“top agar”) method with a top agar prepared as LB agar with only 0.5% w/v agar supplemented with 20 mM MgSO_4_ and 5 mM CaCl_2_^66,67^. Top agar plates were incubated at 37°C for ca 16h before plaque enumeration. High-titer stocks of bacteriophages were generated using the plate overlay method. Briefly, top agar plates were set up to grow almost confluent plaques of a given phage and then covered with 12 mL of SM buffer. After careful agitation for 24-72 hours at 4°C, the suspension on each plate was pipetted off and centrifuged at 8,000 g for 10 minutes. Supernatants were sterilized with few drops of chloroform and stored in the dark at 4°C.

### Bacterial strains and strain construction

All bacterial strains used in this work are listed in Supplementary Table 3. The *P. aeruginosa* phage isolation strain *P. aeruginosa* PAO1 *hsdR17* was generated via the use of the established suicide plasmid pEX18-Tc^68^. All remaining mutants were generated using pFOGG-based suicide plasmids (see below). Plasmids were either electroporated (2.5kV / 25 μF / 400 Ω) or mated into their host using *E. coli* JKE201 as donor strain using standard^69^.

### Plasmid construction

Plasmids were commonly constructed using classical restriction-ligation cloning or the method of Gibson et al. (“Gibson Assembly”)^70^ by ligating PCR products guided by 25 bp overlaps. Point mutations in plasmids were introduced by PCR with partially overlapping primers using the method of Liu and Naismith^71^. *E. coli* strain EC100 *pir(+)* was the host strain of al molecular cloning and successful plasmid construction was routinely assessed by Sanger Sequencing. All oligonucleotide primers used in this study are listed in Supplementary Table 4 and all plasmids are listed in Supplementary Table 5.

### Bacteriophage isolation

Bacteriophages described in this study were isolated between March 2019 and March 2021 generally as described previously^72^ and a complete list of all used phages can be found in Supplementary Table 1. To isolate phages infecting bacteria in stationary phase (see also Fig. 2a), we used 10 ml of deep-dormant culture of *E. coli* K-12 MG1655 or *P. aeruginosa* PAO1 *hsdR17* as described previously and then added 50-300μL of phage precipitate^35^. Upon addition, a 100 μl aliquot was plated by double agar overlay and used to estimate the number of phages initially present. Upon agitation in Erlenmeyer flasks for 48-168h at 37°C, the cultures were centrifuged at full speed for 5 minutes and the supernatants transferred to fresh tubes. Supernatants were sterilized with a few drops of chloroform before 100 μl were plated by double agar overlay and the rest was stored at 4°C. We then counted plaques after overnight incubation at 37°C to evaluate whether phage replication had occurred during the cultivation on the deep-dormant culture. Phage isolates with the ability to replicate on dormant hosts were propagated and stocked as described previously^72^.

### Antibiotic treatment and phage infection assays

Time-resolved kill curves with phages and antibiotics were generally performed as described previously^35^. In addition to determining viable cell counts, we also recorded the free phage titer and the number of infected cells when appropriate. Free phages were sampled as described by Bryan and colleagues and subsequently spotted onto a top agar plate of the corresponding host^14^. The number of infected cells was determined by spotting the samples from the serial dilutions used for the viable cell quantification onto a top agar plate of the respective host bacterium. In the absence of free phages, plaques originate from infected bacteria as centres of infection.

The experiment shown in Fig. 5c was performed by growing a culture of *P. aeruginosa ΔpelΔpsl* into stationary phase for 36h and then diluting it back into fresh medium containing 1mM of DL-serine hydroxamate before incubation for 12h at 37°C shaking. Subsequently, antibiotic and phage treatment were started, viable cells and free phages were sampled and quantified as usual. As control, a parallel experiment (Extended Data Fig. 5e) with a culture freshly diluted 1:10 into fresh medium was performed with the same treatment conditions.

### Bacteriophage genome sequencing, assembly, and annotation

Bacteriophage genomes were purified using the Norgen Biotek Phage DNA Isolation Kit and sequenced at the Microbial Genome Sequencing Center (MiGS) as described previously^72^. Genome assembly and downstream analyses were performed using Geneious Prime 2021.0.1 as described previously^72^. We annotated the genome of Paride and the other newly isolated phages based on a draft annotation generated using multiPhATE^73^ followed by manual curation.

### Sequence alignments and phylogenetic analyses

For the phylogeny shown in Fig. 2d, the major capsid protein, terminase large subunit, and DNA polymerase were extracted from several phages belonging to group 2.2 of jumbo phages^40^ and distantly related myoviruses T4 (GenBank accession NC_000866.4) and Cr30 (GenBank accession NC_025422.1) as outgroup. Besides Paride and its closely related isolates described in this study, we included *Agrobacterium* phage Atu_ph07 (GenBank accession NC_042013.1), *Escherichia* phage PBECO4 (GenBank accession NC_027364.1), *Salmonella* phage Munch (GenBank accession MK268344.1)), and *Xanthomonas* phage XacN1 (GenBank accession AP018399.1). The phylogeny was generated as described previously for other bacteriophages^72^. Briefly, amino acid sequences were aligned using MAFFT v7.450^74^ implemented in Geneious Prime 2021.0.1, manually curated, and then concatenated to calculate a Maximum-Likelihood phylogeny using PhyML 3.3.20180621^75^ implemented in Geneious Prime 2021.0.1.

### Morphological analyses by transmission electron microscopy

The virion morphology of Paride was analyzed by transmission electron microscopy following common procedures in the field^76^. Briefly, 5 μl drops of high-titer lysate were adsorbed to 400 mesh carbon-coated grids, which were rendered hydrophilic using a glow-discharger at low vacuum conditions. They were subsequently stained on 5 μl drops of 2% (w/v) uranyl acetate. Samples were examined using an FEI Tecnai G2 Spirit transmission electron microscope (FEI Company, Hillsboro, Oregon, USA) operating at 80-kV accelerating voltage. Images were recorded with a side-mounted Olympus Veleta CCD camera 4k using EMSIS RADIUS software at a nominal magnification of typically 150,000×.

### Clinical isolate selection and infection

Clinical isolates of *P. aeruginosa* from cystic fibrosis patients were generously shared by the University Hospital of Basel via Prof. Urs Jenal (Supplementary Table 7). Candidates for testing of Paride susceptibility were chosen randomly with preference for high-tolerance isolates described in the study by Santi, Manfredi, and colleagues^77^. We first screened a total of 91 *P. aeruginosa* isolates first for general susceptibility to Paride (with 21/91 being susceptible) and then selected ten isolates for stationary phase infections based on robust growth in M9Rich and LB agar top agars. We determined the MIC of relevant strains as described before^35^ in M9Rich (Supplementary Table 8). These strains were then grown to late stationary phase like in regular Paride infection experiments (see above) and infected with Paride at an MOI of ca. 1:5’000. Free phage titers were determined after 48h of cultivation of 37°C and compared to the inoculum to detect possible phage replication (Fig. 3a).

### Lipopolysaccharides and bacteriophage surface receptors on *P. aeruginosa* PAO1

To gain further insight into the essential host receptor of Paride, we isolated spontaneously resistant mutants by plating bacteria on LB agar plates which had been densely covered with high-titer lysates of the phage. After whole genome sequencing, we determined the efficiency of plating for several phages with different known receptors on these mutants (Supplementary Table 2 and Extended Data Fig. 4). Through the comparison of the EOP, the known structures of different receptor mutants (*ΔwbpL* and *ΔgalU*) and proposed phenotypes for PA5001 (*ssg*) from previous studies, we concluded that the secondary receptor of phage Paride is likely to be at the α-Glucose(III) moiety of the core LPS (Fig. 4). Since the exact structure of the LPS formed by a *P. aeruginosa* PAO1 *ssg* (PA5001) mutant is unknown, we highlighted the sugar suspected to be missing by crossing it off in red (Fig. 4). The remaining residues were represented with dashed lines to indicate that their presence is uncertain. The image is not drawn to scale and was adapted and redrawn from different sources^47-49,63,78,79^.

### Bacterial genome sequencing and assembly

For whole-genome sequencing, genomic DNA was prepared using the GenElute Bacterial Genomic DNA Kit (Sigma-Aldrich, St. Louis, Missouri, USA) according to the manufacturer’s guidelines and sequenced at the Microbial Genome Sequencing Center (MiGS) using the Illumina NextSeq 550 platform. Genome assembly and mutation mapping were performed using *breseq* (https://github.com/barricklab/breseq).

### Experimental adaptation of Paride by passaging on stationary phase cultures

Two parallel cultures of *P. aeruginosa* were grown to stationary phase as described previously^35^ and at the start infected with Paride at an MOI of 1:100’000. Infected cultures (5ml volume) were agitated at 37°C for 72h (first 40 transfers) which was later shortened to 24h (transfers 41 to 71). At each transfer, a sample of each previous infection culture was sterilized with chloroform and diluted 1:100’000 into a freshly grown stationary phase culture. At the end of the experimental evolution, single plaques were picked from both evolutionary lines and used for further experimentation as Paride_1 and Paride_2.

### Quantification of Paride infections using one-step growth curves

One-step growth curve experiments were designed based on established procedures in the field^80,81^. Bacteria were first grown from in M9Glc medium from single colony for 24 hours at 37°C and subsequently diluted back 1:100 for additional 24 hours of cultivation. Fast-growing cultures were generated by an additional 1:100 dilution of this dense culture followed by three hours of cultivation at 37°C shaking. Subsequently, 1 ml of culture was spun down at maximal speed in a tabletop centrifuge and resuspended in 100 μl of fresh M9Glc medium (obtaining ca. 10^9^ CFU/ml). For stationary phase experiments, 1 ml of the original dense culture was used.

Cells with phage at an MOI of ca. 0.1 followed by 15 minutes of adsorption at 37°C shaking before the sample was diluted 1:10’000 into 25 mL of prewarmed medium to prevent further infection cycles. While regularly growing cells were diluted into M9Glc, stationary phase bacteria were diluted back into M9nocarbon, a variant of M9Rich medium where no carbon source and no LB broth are added, to prevent resuscitation when encountering fresh medium. These cultures were agitated in Erlenmeyer flasks using a shaking water bath at 37°C (Julabo SW22). We measured the number of initially infected cells and changes in free phage titers over time by double-layer agar assays as described above.

Latency was determined as the first timepoint where phages could be detected among at least two technical replicates. Burst size was estimated by dividing the average number of free phages at the plateau by the number of infected cells upon dilution.

### Efficiency of plating experiments

The infectivity of a phage on a given host was quantified by determining the efficiency of plating (EOP), i.e., by quantifying its plaque formation on this host in comparison to plaque formation on reference strain *P. aeruginosa* PAO1 *Δpel Δpsl* as described before^72^.

### Quantification and analysis

Quantitative data sets were analysed by calculating mean and standard error of the mean of independent biological replicates for each experiment. Detailed information about replicates and statistical analyses for each experiment is provided in the figure legends. Data were analysed in Microsoft Excel and plotted using R-Studio.

## Acknowledgements

The authors are grateful for helpful discussions with Aisylu Shaidullina, Fabienne Estermann, Dr. Szabolcs Semsey, Prof. Marek Basler, Prof. Martin Loessner, and Prof. Kenn Gerdes. Dr. Mohamed Chami and Carola Alampi of the BioEM lab of the Biozentrum, University of Basel are acknowledged for their support with transmission electron microscopy. Prof. George O’Toole, Prof. Tyler Kokjohn, and the Félix d’Hérelle Reference Center for Bacterial Viruses generously shared bacteriophages used in this work. We thank Dr. Isabella Santi and Dr. Pablo Manfredi for help with strains and plasmids. Fabienne Estermann is acknowledged for her help with isolating phage EMΦ17 and the data analysis. The authors are grateful to ARA Basel and ARA Canius (Lenzerheide) for providing samples of sewage plant inflow. This work was supported by the Swiss National Science Foundation (SNSF) Ambizione Fellowship PZ00P3_180085 and SNSF National Centre of Competence in Research (NCCR) AntiResist.

## Author contributions

A.H. initiated and supervised the study with scholarly advice of U.J.. A.H. and E.M. isolated bacteriophages, performed whole-genome sequencing and phenotypic analyses, and designed most experiments. M.B. generated evolved variants of phage Paride and characterized them. E.M., M.B., and Y.H. performed antibiotic killing and bacteriophage infection experiments. A.E. and U.J. obtained, characterized, and shared clinical isolates of *P. aeruginosa*. A.H. and E.M. wrote the manuscript with input from all authors.

The authors declare no competing interests. Supplementary Information is available for this paper. Correspondence and requests for materials should be addressed to Dr. Alexander Harms.

## Notes

### Competing Interest Statement

The authors have declared no competing interest.

## References

1 Dion, M. B., Oechslin, F. & Moineau, S. Phage diversity, genomics and phylogeny. Nat Rev Microbiol 18, 125–138, doi:10.1038/s41579-019-0311-5 (2020).

2 Chevallereau, A., Pons, B. J., van Houte, S. & Westra, E. R. Interactions between bacterial and phage communities in natural environments. Nat Rev Microbiol 20, 49–62, doi:10.1038/s41579-021-00602-y (2022).

3 Hatfull, G. F., Dedrick, R. M. & Schooley, R. T. Phage Therapy for Antibiotic-Resistant Bacterial Infections. Annu Rev Med, doi:10.1146/annurev-med-080219-122208 (2021).

4 Kortright, K. E., Chan, B. K., Koff, J. L. & Turner, P. E. Phage Therapy: A Renewed Approach to Combat Antibiotic-Resistant Bacteria. Cell Host Microbe 25, 219–232, doi:10.1016/j.chom.2019.01.014 (2019).

5 Murray, C. J. L. et al. Global burden of bacterial antimicrobial resistance in 2019: a systematic analysis. The Lancet, doi:10.1016/S0140-6736(21)02724-0 (2022).

6 Lourenco, M., De Sordi, L. & Debarbieux, L. The Diversity of Bacterial Lifestyles Hampers Bacteriophage Tenacity. Viruses 10, doi:10.3390/v10060327 (2018).

7 Riding, D. Acute Bacillary Dysentery in Khartoum Province, Sudan, with Special Reference to Bacteriophage Treatment: Bacteriological Investigation. J Hyg (Lond) 30, 387–401, doi:10.1017/s0022172400010512 (1930).

8 Brauner, A., Fridman, O., Gefen, O. & Balaban, N. Q. Distinguishing between resistance, tolerance and persistence to antibiotic treatment. Nat Rev Microbiol 14, 320–330, doi:10.1038/nrmicro.2016.34 (2016).

9 Kaldalu, N. et al. In Vitro Studies of Persister Cells. Microbiol Mol Biol Rev 84, doi:10.1128/MMBR.00070-20 (2020).

10 Bergkessel, M., Basta, D. W. & Newman, D. K. The physiology of growth arrest: uniting molecular and environmental microbiology. Nat Rev Microbiol 14, 549–562, doi:10.1038/nrmicro.2016.107 (2016).

11 Verstraete, L., Van den Bergh, B., Verstraeten, N. & Michiels, J. Ecology and evolution of antibiotic persistence. Trends Microbiol, doi:10.1016/j.tim.2021.10.001 (2021).

12 Los, M. et al. Effective inhibition of lytic development of bacteriophages lambda, P1 and T4 by starvation of their host, Escherichia coli. BMC Biotechnol 7, 13, doi:10.1186/1472-6750-7-13 (2007).

13 Pearl, S., Gabay, C., Kishony, R., Oppenheim, A. & Balaban, N. Q. Nongenetic individuality in the host-phage interaction. PLoS Biol 6, e120, doi:10.1371/journal.pbio.0060120 (2008).

14 Bryan, D., El-Shibiny, A., Hobbs, Z., Porter, J. & Kutter, E. M. Bacteriophage T4 Infection of Stationary Phase E. coli: Life after Log from a Phage Perspective. Front Microbiol 7, 1391, doi:10.3389/fmicb.2016.01391 (2016).

15 Ripp, S. & Miller, R. V. Dynamics of the pseudolysogenic response in slowly growing cells of Pseudomonas aeruginosa. Microbiology (Reading) 144 (Pt 8), 2225–2232, doi:10.1099/00221287-144-8-2225 (1998).

16 Los, M. & Wegrzyn, G. Pseudolysogeny. Adv Virus Res 82, 339–349, doi:10.1016/B978-0-12-394621-8.00019-4 (2012).

17 Schooley, R. T. et al. Development and Use of Personalized Bacteriophage-Based Therapeutic Cocktails To Treat a Patient with a Disseminated Resistant Acinetobacter baumannii Infection. Antimicrob Agents Chemother 61, doi:10.1128/AAC.00954-17 (2017).

18 Khatami, A. et al. Bacterial lysis, autophagy and innate immune responses during adjunctive phage therapy in a child. EMBO Mol Med 13, e13936, doi:10.15252/emmm.202113936 (2021).

19 Eskenazi, A. et al. Combination of pre-adapted bacteriophage therapy and antibiotics for treatment of fracture-related infection due to pandrug-resistant Klebsiella pneumoniae. Nat Commun 13, 302, doi:10.1038/s41467-021-27656-z (2022).

20 Livermore, D. M. Has the era of untreatable infections arrived? J Antimicrob Chemother 64 Suppl 1, i29–36, doi:10.1093/jac/dkp255 (2009).

21 Balaban, N. Q. et al. Definitions and guidelines for research on antibiotic persistence. Nat Rev Microbiol 17, 441–448, doi:10.1038/s41579-019-0196-3 (2019).

22 Mizuno, C. M. et al. Isolation and Characterization of Bacteriophages That Infect Citrobacter rodentium, a Model Pathogen for Intestinal Diseases. Viruses 12, doi:10.3390/v12070737 (2020).

23 Golec, P., Karczewska-Golec, J., Los, M. & Wegrzyn, G. Bacteriophage T4 can produce progeny virions in extremely slowly growing Escherichia coli host: comparison of a mathematical model with the experimental data. FEMS Microbiol Lett 351, 156–161, doi:10.1111/1574-6968.12372 (2014).

24 You, L., Suthers, P. F. & Yin, J. Effects of Escherichia coli physiology on growth of phage T7 in vivo and in silico. J Bacteriol 184, 1888–1894, doi:10.1128/JB.184.7.1888-1894.2002 (2002).

25 Nabergoj, D., Modic, P. & Podgornik, A. Effect of bacterial growth rate on bacteriophage population growth rate. Microbiologyopen 7, e00558, doi:10.1002/mbo3.558 (2018).

26 Bokinsky, G. et al. HipA-triggered growth arrest and beta-lactam tolerance in Escherichia coli are mediated by RelA-dependent ppGpp synthesis. J Bacteriol 195, 3173–3182, doi:10.1128/JB.02210-12 (2013).

27 Lopatina, A., Tal, N. & Sorek, R. Abortive Infection: Bacterial Suicide as an Antiviral Immune Strategy. Annu Rev Virol 7, 371–384, doi:10.1146/annurev-virology-011620-040628 (2020).

28 Meeske, A. J., Nakandakari-Higa, S. & Marraffini, L. A. Cas13-induced cellular dormancy prevents the rise of CRISPR-resistant bacteriophage. Nature 570, 241–245, doi:10.1038/s41586-019-1257-5 (2019).

29 Braun, V. FhuA (TonA), the career of a protein. J Bacteriol 191, 3431–3436, doi:10.1128/JB.00106-09 (2009).

30 Silveira, C. B., Luque, A. & Rohwer, F. The landscape of lysogeny across microbial community density, diversity and energetics. Environ Microbiol 23, 4098–4111, doi:10.1111/1462-2920.15640 (2021).

31 Schrader, H. S. et al. Bacteriophage infection and multiplication occur in Pseudomonas aeruginosa starved for 5 years. Can J Microbiol 43, 1157–1163 (1997).

32 Tabib-Salazar, A. et al. T7 phage factor required for managing RpoS in Escherichia coli. Proc Natl Acad Sci U S A 115, E5353–E5362, doi:10.1073/pnas.1800429115 (2018).

33 Harms, A., Fino, C., Sørensen, M. A., Semsey, S. & Gerdes, K. Prophages and Growth Dynamics Confound Experimental Results with Antibiotic-Tolerant Persister Cells. MBio 8, doi:10.1128/mBio.01964-17 (2017).

34 Kragh, K. N. et al. The Inoculation Method Could Impact the Outcome of Microbiological Experiments. Appl Environ Microbiol 84, doi:10.1128/AEM.02264-17 (2018).

35 Maffei, E., Fino, C. & Harms, A. Antibiotic Tolerance and Persistence Studied Throughout Bacterial Growth Phases. Methods Mol Biol 2357, 23–40, doi:10.1007/978-1-0716-1621-5_2 (2021).

36 Hendrix, R. W. Jumbo bacteriophages. Curr Top Microbiol Immunol 328, 229–240, doi:10.1007/978-3-540-68618-7_7 (2009).

37 Drulis-Kawa, Z., Olszak, T., Danis, K., Majkowska-Skrobek, G. & Ackermann, H. W. A giant Pseudomonas phage from Poland. Arch Virol 159, 567–572, doi:10.1007/s00705-013-1844-y (2014).

38 Imam, M. et al. vB_PaeM_MIJ3, a Novel Jumbo Phage Infecting Pseudomonas aeruginosa, Possesses Unusual Genomic Features. Front Microbiol 10, 2772, doi:10.3389/fmicb.2019.02772 (2019).

39 Mendoza, S. D. et al. A bacteriophage nucleus-like compartment shields DNA from CRISPR nucleases. Nature 577, 244–248, doi:10.1038/s41586-019-1786-y (2020).

40 l, M. I., Anantharaman, V., Krishnan, A., Burroughs, A. M. & Aravind, L. Jumbo Phages: A Comparative Genomic Overview of Core Functions and Adaptions for Biological Conflicts. Viruses 13, doi:10.3390/v13010063 (2021).

41 Olszak, T. et al. Pseudomonas aeruginosa PA5oct Jumbo Phage Impacts Planktonic and Biofilm Population and Reduces Its Host Virulence. Viruses 11, doi:10.3390/v11121089 (2019).

42 Wright, R. C. T., Friman, V. P., Smith, M. C. M. & Brockhurst, M. A. Cross-resistance is modular in bacteria-phage interactions. PLoS Biol 16, e2006057, doi:10.1371/journal.pbio.2006057 (2018).

43 Shen, M. et al. Pseudomonas aeruginosa MutL promotes large chromosomal deletions through non-homologous end joining to prevent bacteriophage predation. Nucleic Acids Res 46, 4505–4514, doi:10.1093/nar/gky160 (2018).

44 Le, S. et al. Chromosomal DNA deletion confers phage resistance to Pseudomonas aeruginosa. Sci Rep 4, 4738, doi:10.1038/srep04738 (2014).

45 Pan, X. et al. Genetic Evidence for O-Specific Antigen as Receptor of Pseudomonas aeruginosa Phage K8 and Its Genomic Analysis. Front Microbiol 7, 252, doi:10.3389/fmicb.2016.00252 (2016).

46 Veeranagouda, Y. et al. Ssg, a putative glycosyltransferase, functions in lipo-and exopolysaccharide biosynthesis and cell surface-related properties in Pseudomonas alkylphenolia. FEMS Microbiol Lett 315, 38–45, doi:10.1111/j.1574-6968.2010.02172.x (2011).

47 Choudhury, B., Carlson, R. W. & Goldberg, J. B. The structure of the lipopolysaccharide from a galU mutant of Pseudomonas aeruginosa serogroup-O11. Carbohydr Res 340, 2761–2772, doi:10.1016/j.carres.2005.09.017 (2005).

48 King, J. D., Kocincova, D., Westman, E. L. & Lam, J. S. Review: Lipopolysaccharide biosynthesis in Pseudomonas aeruginosa. Innate Immun 15, 261–312, doi:10.1177/1753425909106436 (2009).

49 Lam, J. S., Taylor, V. L., Islam, S. T., Hao, Y. & Kocincova, D. Genetic and Functional Diversity of Pseudomonas aeruginosa Lipopolysaccharide. Front Microbiol 2, 118, doi:10.3389/fmicb.2011.00118 (2011).

50 Harms, A., Maisonneuve, E. & Gerdes, K. Mechanisms of bacterial persistence during stress and antibiotic exposure. Science 354, doi:10.1126/science.aaf4268 (2016).

51 Hauryliuk, V., Atkinson, G. C., Murakami, K. S., Tenson, T. & Gerdes, K. Recent functional insights into the role of (p)ppGpp in bacterial physiology. Nat Rev Microbiol 13, 298–309, doi:10.1038/nrmicro3448 (2015).

52 Navarro Llorens, J. M., Tormo, A. & Martinez-Garcia, E. Stationary phase in gram-negative bacteria. FEMS Microbiol Rev 34, 476–495, doi:10.1111/j.1574-6976.2010.00213.x (2010).

53 Pletzer, D. et al. The Stringent Stress Response Controls Proteases and Global Regulators under Optimal Growth Conditions in Pseudomonas aeruginosa. mSystems 5, doi:10.1128/mSystems.00495-20 (2020).

54 Gu Liu, C. et al. Phage-Antibiotic Synergy Is Driven by a Unique Combination of Antibacterial Mechanism of Action and Stoichiometry. mBio 11, doi:10.1128/mBio.01462-20 (2020).

55 Los, M., Wegrzyn, G. & Neubauer, P. A role for bacteriophage T4 rI gene function in the control of phage development during pseudolysogeny and in slowly growing host cells. Res Microbiol 154, 547–552, doi:10.1016/S0923-2508(03)00151-7 (2003).

56 Dimitriu, T. et al. Bacteriostatic antibiotics promote CRISPR-Cas adaptive immunity by enabling increased spacer acquisition. Cell Host Microbe 30, 31–40 e35, doi:10.1016/j.chom.2021.11.014 (2022).

57 Gallet, R., Lenormand, T. & Wang, I. N. Phenotypic stochasticity protects lytic bacteriophage populations from extinction during the bacterial stationary phase. Evolution 66, 3485–3494, doi:10.1111/j.1558-5646.2012.01690.x (2012).

58 Cho, H., Uehara, T. & Bernhardt, T. G. Beta-lactam antibiotics induce a lethal malfunctioning of the bacterial cell wall synthesis machinery. Cell 159, 1300–1311, doi:10.1016/j.cell.2014.11.017 (2014).

59 Zambrano, M. M. & Kolter, R. GASPing for life in stationary phase. Cell 86, 181–184, doi:10.1016/s0092-8674(00)80089-6 (1996).

60 Joers, A., Liske, E. & Tenson, T. Dividing subpopulation of Escherichia coli in stationary phase. Res Microbiol 171, 153–157, doi:10.1016/j.resmic.2020.02.002 (2020).

61 Hirsch-Kauffmann, M., Herrlich, P., Ponta, H. & Schweiger, M. Helper function of T7 protein kinase in virus propagation. Nature 255, 508–510, doi:10.1038/255508a0 (1975).

62 Joers, A. et al. Muropeptides Stimulate Growth Resumption from Stationary Phase in Escherichia coli. Sci Rep 9, 18043, doi:10.1038/s41598-019-54646-5 (2019).

63 Meadow, P. M. & Wells, P. L. Receptor Sites for R-type Pyocins and Bacteriophage E79 in the Core Part of the Lipopolysaccharide of Pseudomonas aeruginosa PAC1. Microbiology 108, 339–343, doi:https://doi.org/10.1099/00221287-108-2-339 (1978).

64 Jarrell, K. & Kropinski, A. M. Identification of the cell wall receptor for bacteriophage E79 in Pseudomonas aeruginosa strain PAO. J Virol 23, 461–466, doi:10.1128/JVI.23.3.461-466.1977 (1977).

65 Budzik, J. M., Rosche, W. A., Rietsch, A. & O’Toole, G. A. Isolation and characterization of a generalized transducing phage for Pseudomonas aeruginosa strains PAO1 and PA14. J Bacteriol 186, 3270–3273, doi:10.1128/JB.186.10.3270-3273.2004 (2004).

66 Kauffman, K. M. & Polz, M. F. Streamlining standard bacteriophage methods for higher throughput. MethodsX 5, 159–172, doi:10.1016/j.mex.2018.01.007 (2018).

67 Kropinski, A. M., Mazzocco, A., Waddell, T. E., Lingohr, E. & Johnson, R. P. Enumeration of bacteriophages by double agar overlay plaque assay. Methods Mol Biol 501, 69–76, doi:10.1007/978-1-60327-164-6_7 (2009).

68 Hoang, T. T., Karkhoff-Schweizer, R. R., Kutchma, A. J. & Schweizer, H. P. A broad-host-range Flp-FRT recombination system for site-specific excision of chromosomally-located DNA sequences: application for isolation of unmarked Pseudomonas aeruginosa mutants. Gene 212, 77–86, doi:10.1016/s0378-1119(98)00130-9 (1998).

69 Harms, A. et al. A bacterial toxin-antitoxin module is the origin of inter-bacterial and inter-kingdom effectors of Bartonella. PLoS Genet 13, e1007077, doi:10.1371/journal.pgen.1007077 (2017).

70 Gibson, D. G. et al. Enzymatic assembly of DNA molecules up to several hundred kilobases. Nat Methods 6, 343–345, doi:10.1038/nmeth.1318 (2009).

71 Liu, H. & Naismith, J. H. An efficient one-step site-directed deletion, insertion, single and multiple-site plasmid mutagenesis protocol. BMC Biotechnol 8, 91, doi:10.1186/1472-6750-8-91 (2008).

72 Maffei, E. et al. Systematic exploration of Escherichia coli phage-host interactions with the BASEL phage collection. PLoS Biol 19, e3001424, doi:10.1371/journal.pbio.3001424 (2021).

73 Ecale Zhou, C. L. et al. multiPhATE: bioinformatics pipeline for functional annotation of phage isolates. Bioinformatics 35, 4402–4404, doi:10.1093/bioinformatics/btz258 (2019).

74 Katoh, K. & Standley, D. M. MAFFT multiple sequence alignment software version 7: improvements in performance and usability. Mol Biol Evol 30, 772–780, doi:10.1093/molbev/mst010 (2013).

75 Guindon, S. et al. New algorithms and methods to estimate maximum-likelihood phylogenies: assessing the performance of PhyML 3.0. Syst Biol 59, 307–321, doi:10.1093/sysbio/syq010 (2010).

76 Aziz, R. K., Ackermann, H. W., Petty, N. K. & Kropinski, A. M. Essential Steps in Characterizing Bacteriophages: Biology, Taxonomy, and Genome Analysis. Methods Mol Biol 1681, 197–215, doi:10.1007/978-1-4939-7343-9_15 (2018).

77 Santi, I., Manfredi, P., Maffei, E., Egli, A. & Jenal, U. Evolution of Antibiotic Tolerance Shapes Resistance Development in Chronic Pseudomonas aeruginosa Infections. mBio 12, doi:10.1128/mBio.03482-20 (2021).

78 Carim, S. et al. Systematic discovery of pseudomonad genetic factors involved in sensitivity to tailocins. ISME J 15, 2289–2305, doi:10.1038/s41396-021-00921-1 (2021).

79 Burrows, L. L. Pseudomonas aeruginosa twitching motility: type IV pili in action. Annu Rev Microbiol 66, 493–520, doi:10.1146/annurev-micro-092611-150055 (2012).

80 Kropinski, A. M. Practical Advice on the One-Step Growth Curve. Methods Mol Biol 1681, 41–47, doi:10.1007/978-1-4939-7343-9_3 (2018).

81 Ellis, E. L. & Delbruck, M. The Growth of Bacteriophage. J Gen Physiol 22, 365–384, doi:10.1085/jgp.22.3.365 (1939).

