## Supplementary figures and images for "Phage Paride hijacks bacterial stress responses to kill dormant, antibiotic-tolerant cells"

### Extended Data Fig. 1

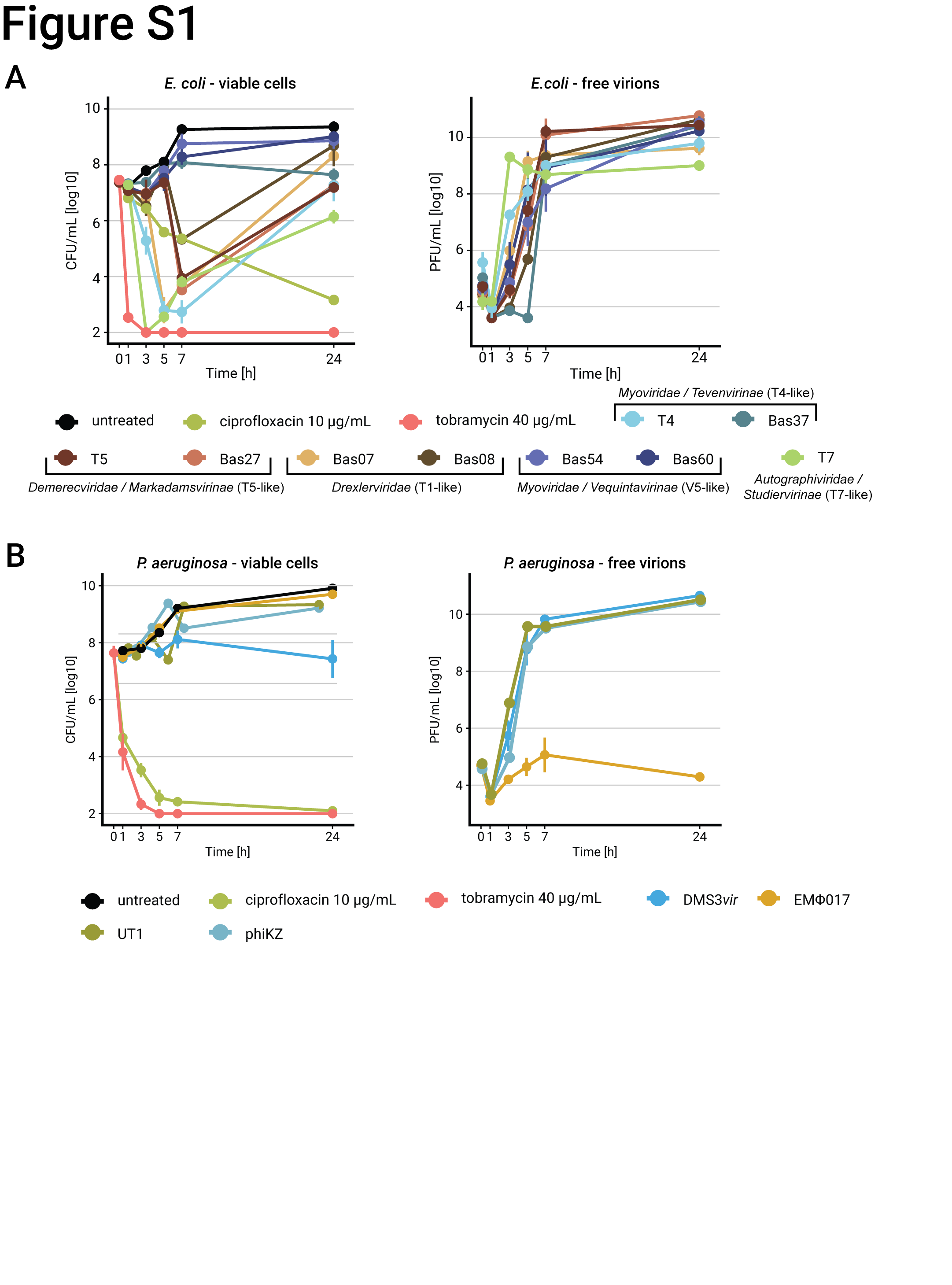

### Extended Data Fig. 2

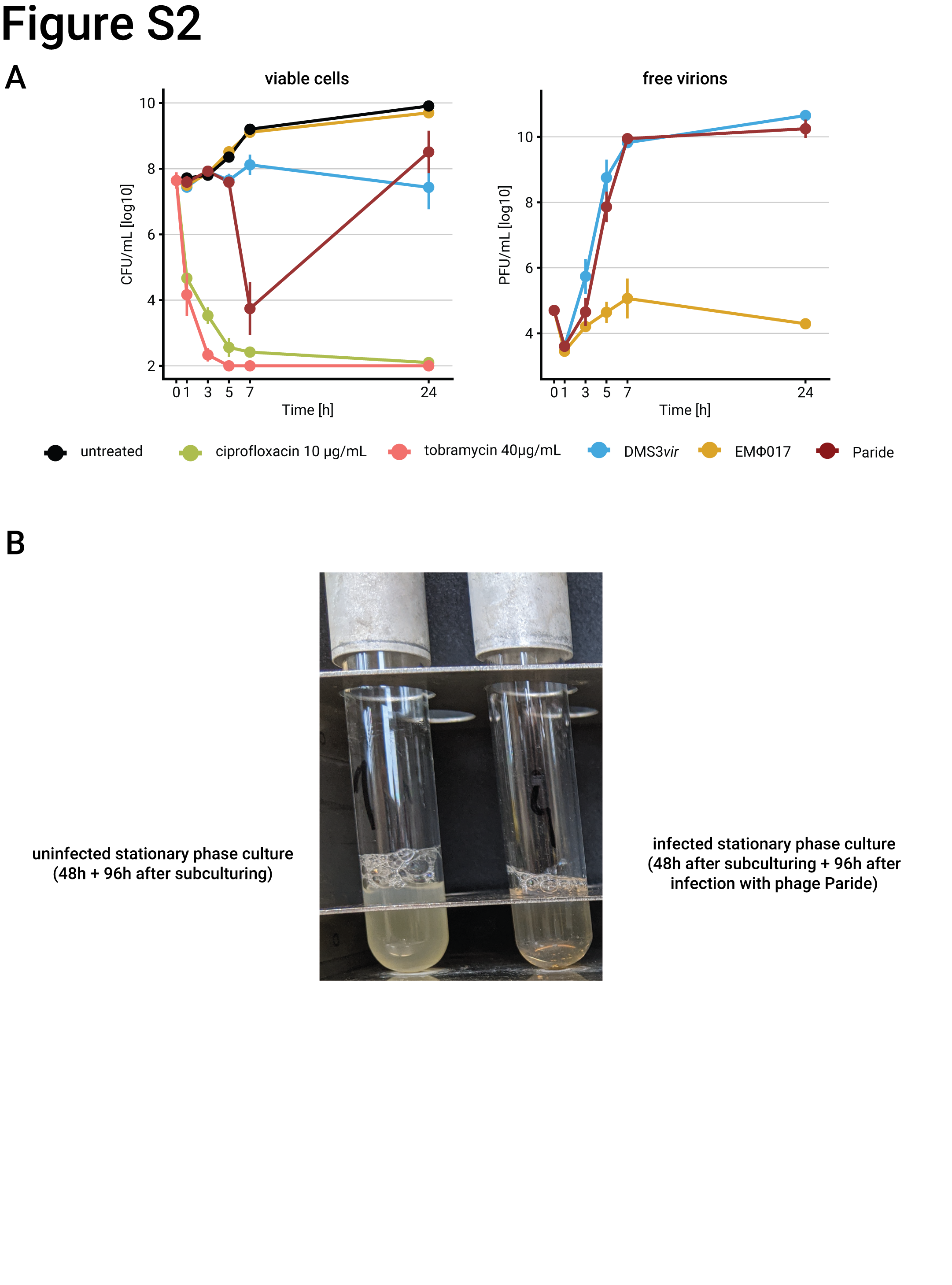

### Extended Data Fig. 3

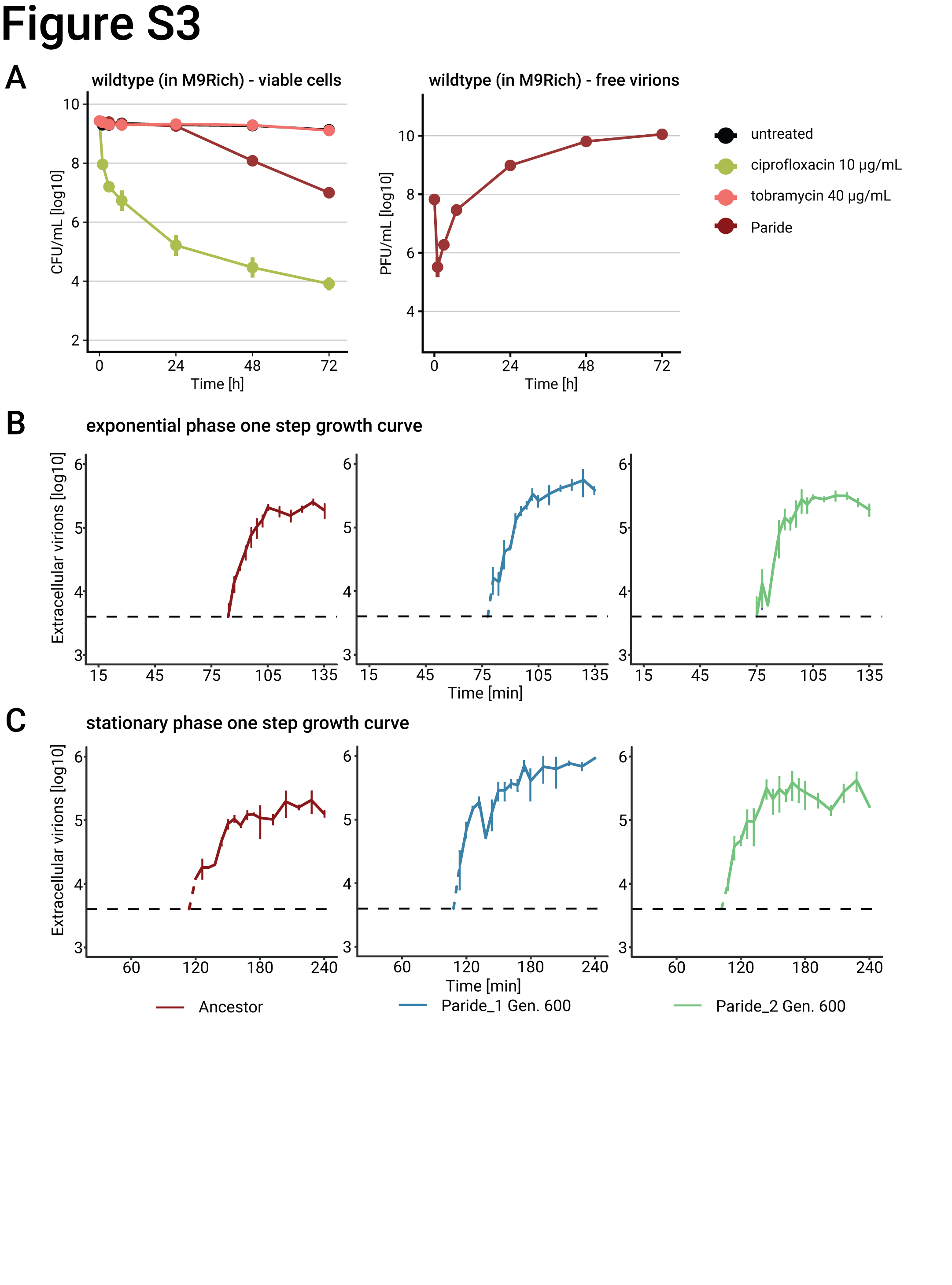

### Extended Data Fig. 4

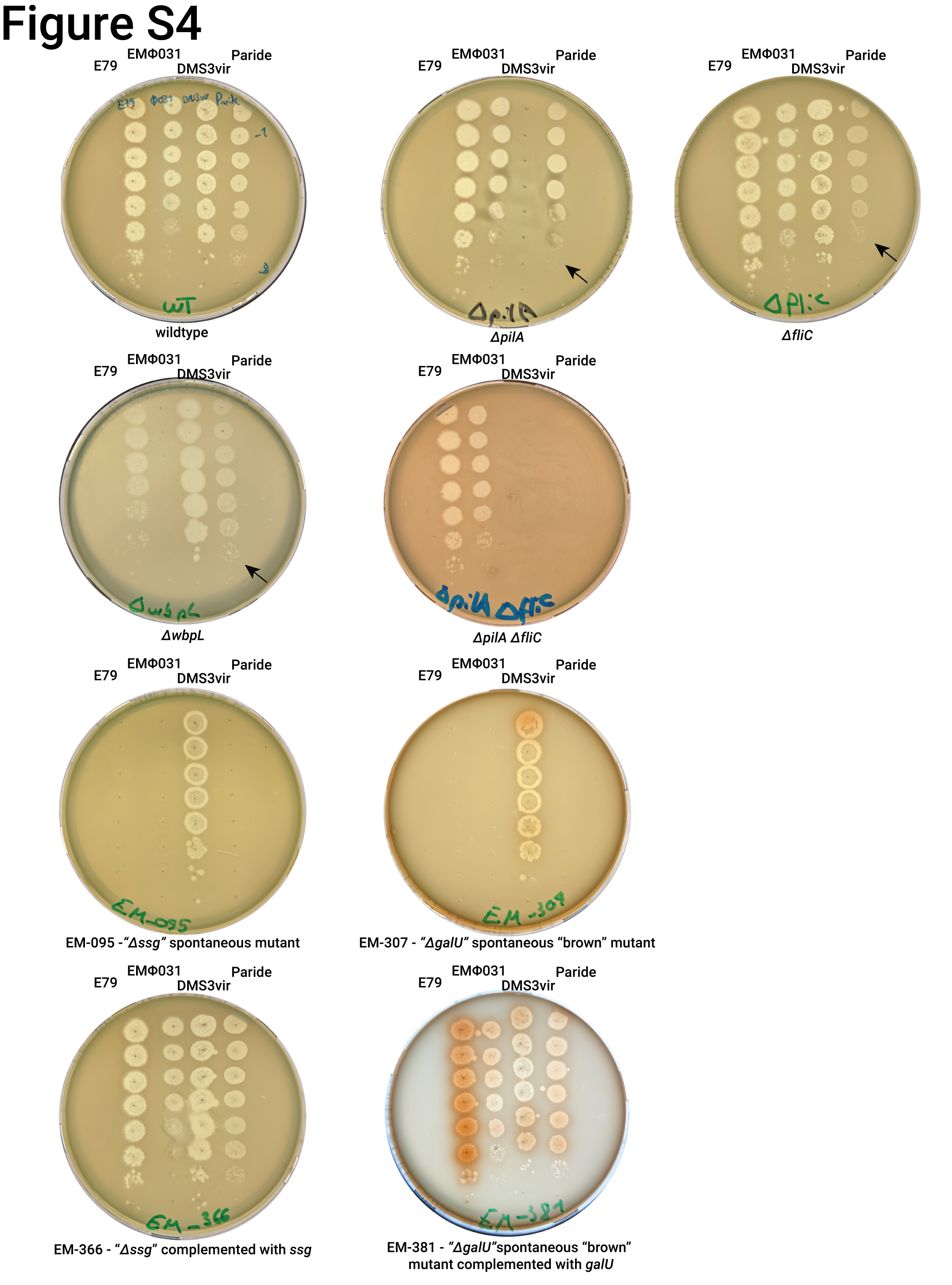

### Extended Data Fig. 5

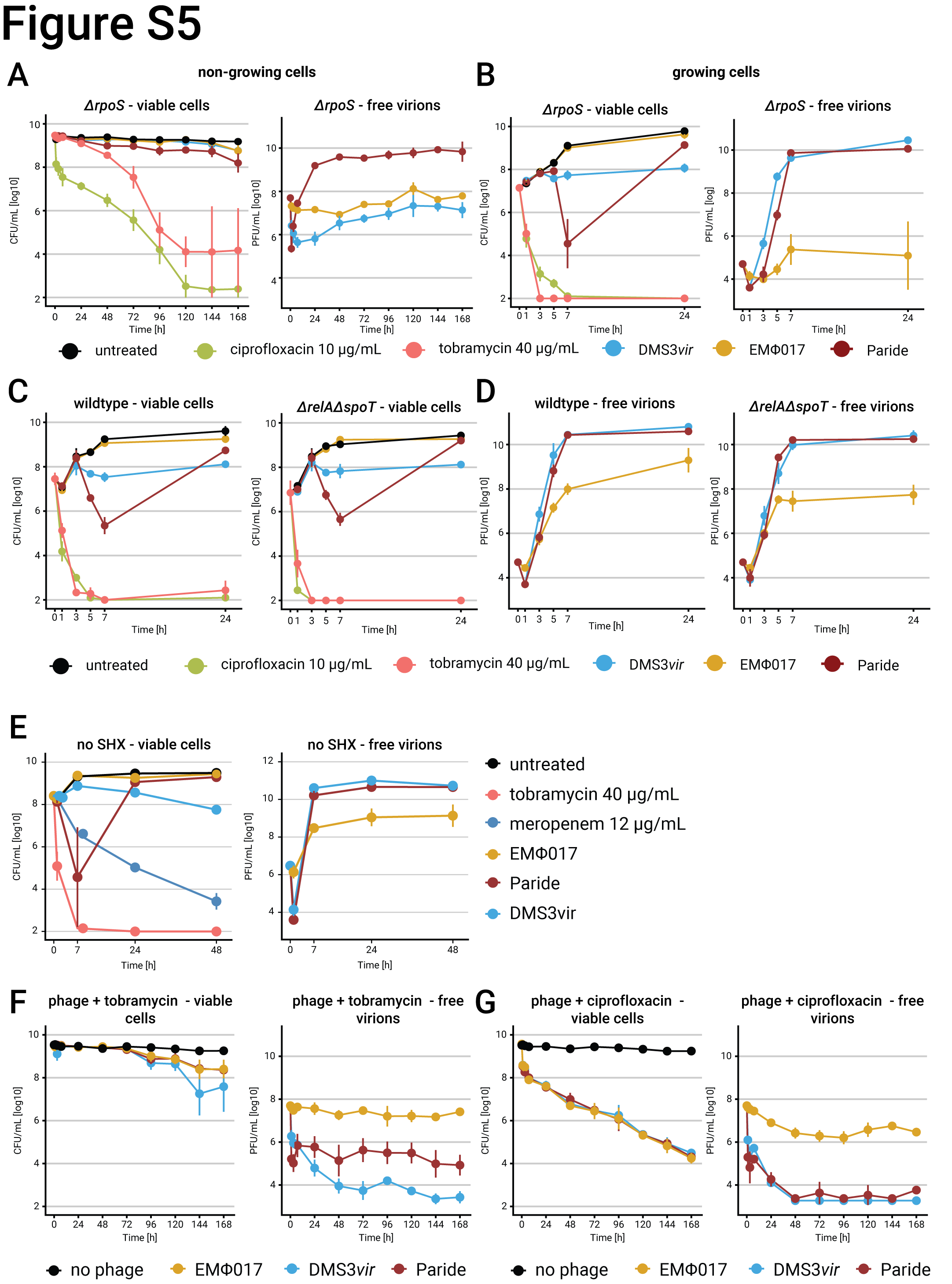
