## Supplementary Table 1 for "Phage Paride hijacks bacterial stress responses to kill dormant, antibiotic-tolerant cells"

### Table S1 – Bacteriophages used in this study

| **Common name** | **Systematic name** | **Host** | **Isolation source** | **Reference** |
| --- | --- | --- | --- | --- |
| T4 | vB_EcoM_T4 | *E. coli* K-12 MG1655 | NA | our laboratory collection |
| Bas37 | vB_EcoM_KarlGJung | *E. coli* K-12 MG1655 | NA | Maffei, Shaidullina, et al., *PLoS Biology* (2021) |
| T5 | vB_EcoS_T5 | *E. coli* K-12 MG1655 | NA | our laboratory collection |
| Bas27 | vB_EcoS_TrudiGerster | *E. coli* K-12 MG1655 | NA | Maffei, Shaidullina, et al., *PLoS Biology* (2021) |
| T7 | vB_EcoP_T7 | *E. coli* K-12 MG1655 | NA | our laboratory collection |
| Bas07 | vB_EcoS_JakobBernoulli | *E. coli* K-12 MG1655 | NA | Maffei, Shaidullina, et al., *PLoS Biology* (2021) |
| Bas08 | vB_EcoS_DanielBernoulli | *E. coli* K-12 MG1655 | NA | Maffei, Shaidullina, et al., *PLoS Biology* (2021) |
| Bas54 | vB_EcoM_MaxBurger | *E. coli* K-12 MG1655 | NA | Maffei, Shaidullina, et al., *PLoS Biology* (2021) |
| Bas60 | vB_EcoM_PaulScherrer | *E. coli* K-12 MG1655 | NA | Maffei, Shaidullina, et al., *PLoS Biology* (2021) |
| DMS3vir | vB_PaeS_DMS3vir | *P. aeruginosa* PAO1 | NA | Budzik et al., *J Bacteriol* (2004) |
| EMΦ017 | vB_PaeS_EMΦ017 | *P. aeruginosa* PAO1 | marsh sample (Sursee, CH; March 2019) | This study |
| EMΦ031 | unknown | *P. aeruginosa* PAO1 | sewage inflow (ARA Canius, Lenzerheide, CH; April 2019) | This study |
| UT1 | vB_PaeM_UT1 | *P. aeruginosa* PAO1 | NA | Schrader et al., *Can J Microbiol* (1997) |
| phiKZ | vB_PaeM_phiKZ | *P. aeruginosa* PAO1 | NA | Krylov and Zhazykov, *Genetika* (1978) |
| Paride | vB_PaeM_Paride | *P. aeruginosa* PAO1 | rotting plant material from Hörnli cemetery (Basel, CH; May 2019) | This study |
| Ettore | vB_PaeM_Ettore | *P. aeruginosa* PAO1 | sewage inflow (ARA Basel, CH; March 2020) | This study |
| Deiofobo | vB_PaeM_Deiofobo | *P. aeruginosa* PAO1 | sewage inflow (ARA Basel, CH; October 2020) | This study |
| Cassandra | vB_PaeM_Cassandra | *P. aeruginosa* PAO1 | sewage inflow (ARA Basel, CH; February 2021) | This study |
