## Supplementary Table 2 for "Phage Paride hijacks bacterial stress responses to kill dormant, antibiotic-tolerant cells"

### Table S2 – Phage susceptibility of *P. aeruginosa* PAO1 surface receptor mutants

(+) – clear plaques, (+/-) opaque plaque, (-) reduced efficiency of plating, (--) loss of lytic activity

| **Strain** | **Phenotype** | **E79 activity** | **EM****Φ031**  **activity** | **DMS3vir activity** | **Paride activity** |
| --- | --- | --- | --- | --- | --- |
| PAO1 | PAO1 wildtype strain | (+) | (+) | (+) | (+) |
| PAO1 *ΔpelΔpsl* | Wildtype strain of this study | (+) | (+) | (+) | (+) |
| PAO1 *ΔpilA* | Lack of type IV pili | (+) | (+) | (--) | (+/-) |
| PAO1 *ΔfliC* | Lack of flagella | (+) | (+) | (+) | (+/-) |
| PAO1 *ΔpelΔpsl ΔpilA ΔfliC* | Lack of type IV pili and lack of flagella | (+) | (+) | (--) | (--) |
| PAO1 *ΔpelΔpsl ΔwpbL* | Lack of glucosyltransferase WbpL essential for the  initiation of O-antigen synthesis | (+) | (--) | (+) | (+/-) |
| PAO1 *ΔpelΔpsl ΔgalU* | Lack of uridylyltransferase GalU essential for the synthesis of LPS and O-antigen precursors | (--) | (--) | (+) | (--) |
| PAO1 *ΔpelΔpsl Δssg* | Lack of glucosyltransferase Ssg (PA5001) essential for the  synthesis of O-antigen | (--) | (--) | (+) | (--) |
| PAO1 *ΔpelΔpsl* “EM-095” | Spontaneous mutant resistant to Paride with a(GGC 🡪 G‑C deletion at position 5’618’713 that causes a frameshift in *ssg* (PA5001) | (--) | (--) | (+) | (--) |
| PAO1 *ΔpelΔpsl* “EM-307” | Spontaneous mutant with “brown” phenotype resistant to Paride; carries a deletion of 44’015bp between positions 2’198’124-2’230’850 and 2’231’737-2’242’134 encompassing *galU* | (--) | (--) | (+) | (--) |
| PAO1 *ΔpelΔpsl* “EM-366” | EM-095 complemented with SC101miniTn7-Gm-Plac_PA5001 (*ssg*) | (+) | (+) | (+) | (+) |
| PAO1 *ΔpelΔpsl* “EM-381” | EM-307 complemented with SC101miniTn7-Gm-Plac_*galU* | (+) | (+) | (+) | (+) |

Phage E79 is know to depend on structures in the LPS core of *P. aeruginosa* PAO1 (Meadow and Wells, *Microbiology* (1978)) while phage DMS3*vir* targets type IV pili (Budzik et al., *J Bacteriol* (2004)). Newly isolated phage EMΦ031 requires expression of the LPS O-antigen of *P. aeruginosa* PAO1. Base pair coordinates refer to the *P. aeruginosa* PAO1 reference genome (NCBI GenBank accession AE004091).
