## Supplementary Table 3 for "Phage Paride hijacks bacterial stress responses to kill dormant, antibiotic-tolerant cells"

### Table S3 - Strains used in this study

| **Strains** | **Description** | **Reference** |
| --- | --- | --- |
| *E. coli* K-12 MG1655 | Wildtype strain of *E. coli* | our laboratory collection |
| *P. aeruginosa* PAO1 | Parental strain of all *P. aeruginosa* PAO1 strains used in this study | our laboratory collection |
| PAO1 *hsdR17* | Phage isolation strain of *P. aeruginosa* PAO1 where the *hsdR* component of the *hsdRMS* system was inactivated | this study |
| PAO1 *ΔpelΔpsl* | Strain with reduced biofilm formation, used as wildtype strain for all experiments unless stated otherwise | Broder et al., *Nat Microbiol* (2016) |
| PAO1 *ΔpelΔpsl ΔrelAΔspoT* | Strain lacking the (p)ppGpp producing and degrading enzymes RelA and SpoT | this study |
| PAO1 *ΔpelΔpsl ΔrpoS* | Strain lacking the RpoS sigma factor | this study |
| PAO1 *ΔpelΔpsl ΔwbpL* | Strain lacking the glycosyltransferase WbpL essential for O‑antigen biosynthesis initiation | this study |
| PAO1 *ΔpelΔpsl* *ΔgalU* | Strain lacking the uridylyltransferase GalU essential for the biosynthesis of LPS and O-antigen precursors | this study |
| PAO1 *ΔpilA* | Strain lacking the major pilin PilA, unable to assemble type IV pili | Laventie et al., *Cell Host Microbe* (2019) |
| PAO1 *ΔfliC* | Strain lacking the flagellin FliC, unable to assemble flagella | Laventie et al., *Cell Host Microbe* (2019) |
| PAO1 *ΔpelΔpsl* *ΔpilA ΔfliC* | Strain lacking PilA and FliC, unable to assemble pili and flagella | Prof. Urs Jenal |
| PAO1 *ΔpelΔpsl* “EM-095” | Spontaneous mutant resistant to Paride carrying a point deletion at position 5’618’713 (GGC 🡪G-C) in the PA5001 gene causing a frameshift in the gene. | This study |
| PAO1 *ΔpelΔpsl* “EM-307” | Spontaneous mutant with “brown” phenotype resistant to Paride carrying a deletion of 44’015bp between positions 2’198’124-2’230’850 and 2’231’737-2’242’134 around *galU* | This study |
| PAO1 *ΔpelΔpsl* “EM-366” | EM-095 complemented with SC101miniTn7-Gm-Plac_PA5001/ssg | This study |
| PAO1 *ΔpelΔpsl* “EM-381” | EM-307 complemented with SC101miniTn7-Gm-Plac_*galU* | This study |

Base pair coordinates refer to the *P. aeruginosa* PAO1 reference genome (NCBI GenBank accession AE004091).
