## Supplementary Table 4 for "Phage Paride hijacks bacterial stress responses to kill dormant, antibiotic-tolerant cells"

### Table S4 – Oligonucleotide primers used in this study

| **Primers** | **Description** | **Sequence 5’->3’** |
| --- | --- | --- |
| prAH2057 | Linearisation of pFOGG vector | CATAAAGTGTAAAGCCTGGGG |
| prAH2058 | Linearisation of pFOGG vector | GCTATCTGGACAAGGGAAAAC |
| prAH2079 | Shuttling primers from pEX18-Tc to pFOGG | TTGCGTTTTCCCTTGTCCAGATAGCCCAGTCACGACGTTGTAAAAC |
| prAH2283 | Shuttling primers from pEX18-Tc to pFOGG | GGCACCCCAGGCTTTACACTTTATGCAGGAAACAGCTATGACCATG |
| prEM0028 | To amplify *hsdR* insert | CGCCAT**TCTAGA**CTCATCGAAGCCGGTGACGAATT |
| prEM0029 | To amplify *hsdR* insert | CGCCAG**GAGCTC**AACTCGACAATAAGCCGGGCAAG |
| prEM0005 | To introduce M353(ATG->TGA) stop mutation in *hsdR* sequence | TGCGGTCACAAAGGGTCGGCTACGCTG |
| prEM0006 | To introduce M353(ATG->TGA) stop mutation in *hsdR* sequence | CTTTGTGACCGCAGTTCGATTCCATCATC |
| prEM0013 | Verification of *hsdR17* mutation | ATCGCTGGCCAACATCATCG |
| prEM0014 | Verification of *hsdR17* mutation | GTGCGCCTCGTCGATCAATA |
| prEM0135 | Verification of *relA* deletion | GTGACTGGCAACTGACTCTGG |
| prEM0136 | Verification of *relA* deletion | GATCGACCTTGAGATGCCG |
| prEM0070 | Verification of *spoT* deletion | GTCTTCGGCAACCTCTACGGCA |
| prEM0071 | Verification of *spoT* deletion | GTCGTCGCCATAGAAGGCAACC |
| prEM0072 | Verification of *rpoS* deletion | AGTTAGTACGTCGGTACCTGC |
| prEM0073 | Verification of *rpoS* deletion | AACATCACCGAGAAGAAGGA |
| prEM0312 | To amplify *galU* deletion insert | **TTGCGTTTTCCCTTGTCCAGATAGC**CTACTCCTGGATCAGATGCGT |
| prEM0313 | To amplify *galU* deletion insert | **CTCATGATCAAG**AAGGCTCACTGAGCCTCGCC |
| prEM0314 | To amplify *galU* deletion insert | **CCTTCTTGATCA**TGAGAATCCTTCGACA |
| prEM0315 | To amplify *galU* deletion insert | **GGCACCCCAGGCTTTACACTTTATG**TCGCCACCAAGATTTCCTTCA |
| prEM0316 | Verification of *galU* deletion | CGATGACGAACAGATCGAAATC |
| prEM0317 | Verification of *galU* deletion | AGTTCAGCAAGTACGCGGC |
| prEM0059 | To amplify *wbpL* deletion insert | **TTGCGTTTTCCCTTGTCCAGATAGC**GTTCCACCAGTTGGGCTTG |
| prEM0060 | To amplify *wbpL* deletion insert | CGGGTTCCTTGGAAAAATCC |
| prEM0061 | To amplify *wbpL* deletion insert | **GGCACCCCAGGCTTTACACTTTATG**CGACCGCCTTTGATCTATGC |
| prEM0062 | To amplify *wbpL* deletion insert | **GCTTAGGATTTTTCCAAGGAACCCG**CGCGATCATCCAGATCATCAT |
| prEM0066 | Verification of *wbpL* deletion | CTTGTAGTTGATCTGCACACC |
| prEM0067 | Verification of *wbpL* deletion | CTATGCCATCTCCAAATACG |
| prEM0306 | To amplify PA5001 deletion insert | **TTGCGTTTTCCCTTGTCCAGATAGC**TTCGATATAGACATCTTCCAGGC |
| prEM0307 | To amplify PA5001 deletion insert | **CTGATGAAAGTT**TTTACCTGAGAGTAGCAGCCG |
| prEM0308 | To amplify PA5001 deletion insert | **CTCTCAGGTAAA**AACTTTCATCAGACCTTCATCCTC |
| prEM0309 | To amplify PA5001 deletion insert | **GGCACCCCAGGCTTTACACTTTATG**AACTGGGTTATTTCTGCCTGC |
| prEM0310 | Verification of PA5001 deletion | TTCTTGTACGTATTGGTGGGAT |
| prEM0311 | Verification of PA5001 deletion | GAGATCGAAACCGAGCATTAC |
| prEM0305 | To amplify PA5001 region for complementation | **GGCGTTACCCAACTTAATCGCCTTG**AAGAATGCTCGCCGTTGCT |
| prEM0306 | To amplify PA5001 region for complementation | **ATCCGCCAAAACAGGGAATTTATGC**CTTTTCGGCTGCTACTCTCAGGT |
| prEM0318 | To amplify *galU* region for complementation | GGCGTTACCCAACTTAATCGCCTTGGTGACCGAGTGGAAACAGTTCC |
| prEM0319 | To amplify *galU* region for complementation | ATCCGCCAAAACAGGGAATTTATGCGTTCGCCCCATACGAAAAACG |
| prAH2376 | To linearise SC101-miniTn7-Gm-P*lac*-*QI* | GCATAAATTCCCTGTTTTGG |
| prAH2319 | To linearise SC101-miniTn7-Gm-P*lac*-*QI* | CAAGGCGATTAAGTTGGGTAA |
| prAH2381 | To amplify pUC18-miniTn7-Gm for *ori* exchange with SC101 | CACGTTAAGGGATTTTGGTC |
| prAH2382 | To amplify pUC18-miniTn7-Gm for *ori* exchange with SC101 | GGGTCATTATAGCGATTTTTTC |
| prAH2379 | To amplify SC101 *ori* to insert in linearized pUC18-miniTn7-Gm | ACCGAAAAAATCGCTATAATGACCCCTCCTGTTGATAGATCCAGTAATG |
| prAH2380 | To amplify SC101 *ori* to insert in linearized pUC18-miniTn7-Gm | CTCATGACCAAAATCCCTTAACGTGCCGCTGTAACAAGTTGTCTC |
| prAH2391: | To linearise SC101-miniTn7-Gm for insertion of P*lac-lacIQ* | GCACCCCAGGCTTTACACTTTATGCCTGAGTAGGACAAATCCGCC |
| prAH2371 | To linearise SC101-miniTn7-Gm for insertion of P*lac-lacIQ* | ACAGGAAGCAAAGCTGAAAGGAATCCGTTTAAGGGCACCAATAACTG |
| prAH2201 | To exchange ATG to GTG of *sce-I* gene of pFOG | AGAGAAAAGTGAAGTGCATCAAAAAAACCAGGTA |
| prAH2374 | To amplify P*lac-lacIQ* from pAH186SC101_e | GATTCCTTTCAGCTTTGCTTC |
| prAH2375 | To amplify P*lac-lacIQ* from pAH186SC101_e | GCATAAAGTGTAAAGCCTGGG |
| prAH2202 | To exchange ATG to GTG of *sce-I* gene of pFOG | TGCACTTCACTTTTCTCTATCACTGATAGG |
