## Supplementary Table 5 for "Phage Paride hijacks bacterial stress responses to kill dormant, antibiotic-tolerant cells"

### Table S5 - Plasmids used in this study

| **Plasmid** | **Description** | **Reference** |
| --- | --- | --- |
| pEX18-Tc | suicide vector for allelic exchange in *P. aeruginosa* | Hoang et al., *Gene* (1998) |
| pEX18-Tc_*hsdR17* | suicide vector used to generate *hsdR17* mutation to inactivate *hsdR* of PAO1 | this study |
| pFOG | suicide vector for allelic exchange in *P. aeruginosa* | Cianfanelli et al., *BMC Microbiol* (2020) |
| pFOGG_*ΔrelA* | suicide vector used to generate *relA* deletion in PAO1 | this study |
| pFOGG_*ΔspoT* | suicide vector used to generate *spoT* deletion in PAO1 | this study |
| pFOGG_*ΔrpoS* | suicide vector used to generate *rpoS* deletion in PAO1 | this study |
| pFOGG_*ΔgalU* | suicide vector used to generate *galU* deletion in PAO1 | this study |
| pFOGG_*ΔwbpL* | suicide vector used to generate *wbpL* deletion in PAO1 | this study |
| pFOGG_PA5001/*ssg* | suicide vector used to generate *ssg* deletion in PAO1 | this study |
| SC101miniTn7-Gm-Plac_*galU* | complementation of *galU* mutants | this study |
| SC101miniTn7-Gm-Plac_PA5001 | complementation of PA5001 mutant | this study |
| pTNS2 | Helper plasmid for transformation of miniTn7 based plasmids | Choi et al., *Nat Methods* (2005) |
| pFLP2 | FRT/FLP excising plasmid for removal of antibiotic cassette of miniTn7 based constructs | Hoang et al., *Gene* (1998) |
