## Supplementary Table 8 for "Phage Paride hijacks bacterial stress responses to kill dormant, antibiotic-tolerant cells"

### Table S8 - MIC of clinical isolates

Table S8. **MIC values of P. aeruginosa PAO1 Δpel Δpsl, CI249 and CI282 in M9Rich medium**. The values reported are the average of two independent experiments.

| Strain | Tobramycin sulfate MIC [μg/mL] | Ciprofloxacin MIC [μg/mL] |
| --- | --- | --- |
| PAO1 *ΔpelΔpsl* | 1 | 0.125 |
| CI249 | 2 | 0.25 |
| CI282 | >32 | >4 |
